# Circadian Clock Programming of Anticipatory Antiviral Immunity Gates Enteric Virus Infection Susceptibility

**DOI:** 10.64898/2026.05.15.725500

**Authors:** Temitope O. Oshinowo, Robert W. Maples, Mikal A. Woods Acevedo, Broc T. McCune, Hunter Dalton, Ishea Johnson, Devin A. Simpkins, Urbashi Basu, Chaitanya Dende, Vera L. Tarakanova, Julie K. Pfeiffer, John F. Brooks

## Abstract

Susceptibility to viral infection varies widely but is not fully explained by genetics, immune status, or exposure level. We show that time of day strongly influences infection outcome, with up to 100-fold differences in enteric viral burden depending on infection timing. This temporal gating is abolished in mice lacking a functional circadian clock. We identify the antiviral transcription factor IRF1 as a direct target of the circadian transcription factor BMAL1, resulting in rhythmic expression of a basal antiviral gene program prior to infection. Loss of IRF1 eliminates this program and abrogates time-of-day–dependent differences in viral replication. This circuit operates within intestinal myeloid cells, establishing a preexisting antiviral state. These findings indicate that the circadian clock programs host susceptibility in the intestine, before infection occurs.

## INTRODUCTION

Susceptibility to viral infection varies widely among individuals, yet the determinants of this variability remain incompletely understood. Host genetics, immune status, and route/level of exposure all shape infection outcomes, but these factors alone do not fully account for observed differences in viral burden. One fundamental dimension of host biology that has received little attention is timing of infection during the day-night cycle.

The circadian clock is an evolutionarily conserved system that synchronizes physiology with the day-night cycle. By coordinating gene expression, metabolism, and behavior (*1, 2*) the clock enables organisms to anticipate predictable environmental changes rather than merely react to them. Immunologically, circadian regulation influences leukocyte trafficking (*3–6*), cytokine production (*3*), and inflammatory responses (*7–11*). These studies have largely focused on how the clock modulates the magnitude or timing of immune responses after stimulation. Prior studies have shown that circadian timing can influence infection with respiratory and systemic viruses (*12–16*). In these settings, time-of-day effects have been attributed largely to variation in host responses after infection or to virus–host interactions during systemic spread. Whether circadian programs instead establish a preexisting antiviral state that determines susceptibility, particularly in the intestine, remains unclear.

Enteric viral infections provide a powerful context for examining temporal regulation of host defense. In addition to rhythmic food intake, the intestinal mucosa is constantly exposed to environmental antigens and microbial products, requiring strategies that balance nutrient acquisition, metabolism, and immune protection vs. damage (*17*). Mammalian enteric virus infections are common, and outcomes range from mild symptoms to fatal diarrhea or systemic disease. Innate antiviral defenses in this setting rely on basal expression of viral sensors and restriction factors that act early in infection (*18, 19*), often before robust interferon amplification occurs. This raises the possibility that temporal regulation at mucosal surfaces may operate through anticipatory immune programs distinct from those described in systemic or respiratory infections. How these basal antiviral programs are regulated in time, and whether they influence infection outcome, is not well understood.

Here, we investigated whether host susceptibility to enteric viral infection is temporally regulated across the day-night cycle. In mice perorally inoculated with the model enteric picornavirus coxsackievirus B3 (CVB3), we show that infection outcome depends strongly on the time of exposure, revealing daily windows of host resistance and susceptibility. We identify a circadian clock-controlled immune program that operates within myeloid cells of the intestinal lamina propria to regulate IRF1 expression and downstream antiviral gene expression. Disruption of this program abolishes temporal differences in viral susceptibility without altering circadian behavior or feeding rhythms. Together, these findings demonstrate that the circadian clock does not merely tune immune responses after infection but instead establishes a temporally gated and anticipatory antiviral state-a “head start” on innate responses during the rest phase in nocturnal animals. By defining time as a critical axis of host susceptibility, this work reframes clock regulation as a fundamental component of anticipatory antiviral defense.

## RESULTS

### Time of infection determines enteric virus replication efficiency

To test whether host susceptibility to enteric viral infection varies by time of day, we perorally infected mice with coxsackievirus B3 (CVB3) at 4-hour intervals over a 24-hour period beginning at zeitgeber time (ZT) 0 (6 am, lights on) (Fig. 1A). Viral loads were quantified by plaque assays 24 hours after infection. Viral loads exhibited a pronounced dependence on infection timing: mice infected during the early light phase (ZT0 and ZT4) had significantly lower viral titers compared to mice infected during the dark phase, which begins at ZT12 (lights off). This temporal difference was evident throughout the gastrointestinal tract, with viral titers in the small intestine, cecum, and stool exhibiting diurnal oscillations (p < 0.0001) that varied by up to 116-fold depending on infection time (Fig. 1B, Fig. S1A, B). Given that enteric viruses spread by the fecal-oral route, elevated viral titers in stool may influence transmission efficiency.

**Fig. 1.**
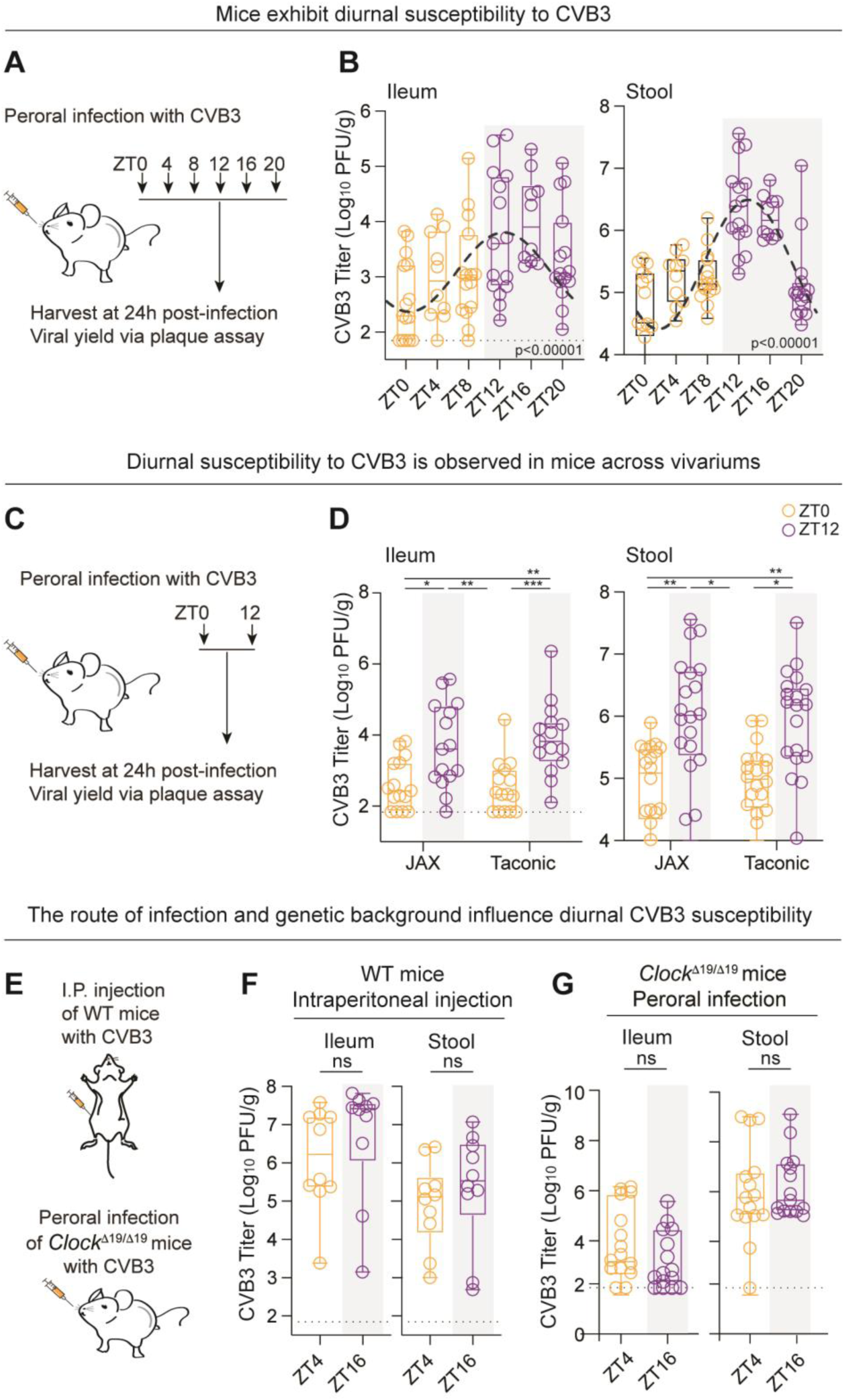
Time of day determines host susceptibility to enteric viral infection. **(A)** Mice were perorally infected with 10^9^ PFU CVB3 at six timepoints across the day-night cycle and tissues were harvested at 24h post-infection followed by plaque assay using HeLa cells. **(B)** Viral load in the ileum and stool of wildtype mice at 24h post-infection. Black dashed line shows rhythmicity analysis (P < 0.0001) of peak and trough within the data set, and dotted line indicates limit of detection. **(C)** Wildtype C57BL/6 mice from The Jackson Laboratory (JAX) versus Taconic Biosciences were perorally infected with 10^9^ PFU CVB3 at one timepoint during the day (ZT0) and one timepoint during the evening (ZT12), followed by viral titer analysis at 24h post-infection. **(D)** Viral load in the ileum and stool of Jackson (JAX) vs. Taconic mice at 24h post-infection. **(E)** Wildtype C57BL/6 mice were intraperitoneally injected with 10^5^ PFU CVB3 at ZT4 or 16 (top) or mice with a disrupted circadian clock (*Clock^Δ19/Δ19^)* were perorally infected with 10^9^ PFU CVB3 at ZT4 or 16 (bottom), and viral load was determined by plaque assay at 24h post-infection. **(F)** Viral load in the ileum and stool of intraperitoneally injected wildtype mice at 24h post-infection. **(G)** Viral load in the ileum and stool of perorally infected *Clock^Δ19/Δ19^* mice at 24h post-infection. PFU, plaque forming units; CVB3, coxsackievirus B3; ZT4, zeitgeber time 4 (10:00 AM); ZT16, zeitgeber time 16 (10:00 PM), *Clock*, circadian locomotor output cycles kaput. Means ± SEM are plotted; *P < 0.05, **P < 0.01; ***P < 0.001; ****P < 0.0001 as determined by Student’s t-test or ANOVA. ns, not significant. Black dashed line indicates rhythmicity ****P <0.0001, generated with CircaCompare.

Next, we examined several factors that may contribute to diurnal variation of infection efficiency: Intestinal microbiota, inoculation route, and a host transcription factor that establishes rhythmic gene expression. First, we determined whether variation in intestinal microbiota influences diurnal effects on CVB3 susceptibility. Our past work demonstrated that microbiota promote enteric virus infection (*20–22*), indicating that microbial communities can strongly influence viral outcomes in the gut. In addition, specific taxa, including segmented filamentous bacteria (SFB) that vary between animal facilities, diurnally modulate host immune pathways and protect against select enteric viruses (*17, 23*). Therefore, we sought to determine whether the observed temporal variation in CVB3 susceptibility was robust across distinct microbiota compositions. To address this, we compared infection in mice obtained from two commercial vendors that harbor distinct and well-characterized microbial communities, including the presence of SFB (Fig. 1C). Time-of-day-dependent differences in CVB3 replication efficiency were preserved in mice across vendors (Fig. 1D, Fig. S1C), suggesting that temporal variation in viral load is largely independent of microbial composition (*17*). Second, we determined whether diurnal effects on CVB3 replication were specific to the natural oral infection route (Fig. 1E). Mice infected via intraperitoneal injection that bypasses the intestinal tract failed to exhibit time-of-day–dependent differences in viral titers (Fig. 1F, Fig. S1D), indicating that temporal gating of viral susceptibility is specific to mucosal infection and is not observed following systemic inoculation. Finally, we examined if the core circadian transcription factor Circadian Locomotor Output Cycles Protein Kaput (CLOCK) influences viral loads across the day-night cycle (Fig. 1E). Mice with a dominant-negative mutation due to the deletion of the CLOCK exon 19, *Clock^Δ19/Δ19^* mice, no longer exhibited time-of-day–dependent differences in viral load following peroral CVB3 infection (Fig. 1G, Fig. S1E). Together, these data demonstrate that host permissiveness to enteric viral infection is temporally regulated, conserved across distinct microbiomes, specific to the oral infection route, and requires an intact circadian clock.

### The circadian clock establishes rhythmic expression of the antiviral regulator IRF1

To identify host immune regulators that could mediate time-of-day–dependent susceptibility, we examined the expression of key transcription factors involved in antiviral immunity at early time points after infection. Among multiple candidates, including *Stat1*, *Stat2*, and several interferon regulatory factors (IRFs), only *Irf1* exhibited robust circadian oscillation in expression (Fig. 2A, Fig. S2A-B). IRF1 is a central transcriptional regulator of antiviral immunity (*24, 25*). Unlike inducible interferon responses that are rapidly engaged following infection, IRF1 contributes to basal expression of antiviral genes, including sensors and effectors that restrict viral replication at initial stages (*26, 27*). IRF1 activity restricts diverse viral pathogens (*28*), yet how its expression is regulated under homeostatic conditions *in vivo* remains poorly defined. We found that *Irf1* expression peaked during the light phase and reached its nadir during the dark phase, coinciding with periods of maximal and minimal host resistance, respectively (Fig. 2A, Fig. 1B).

**Fig. 2.**
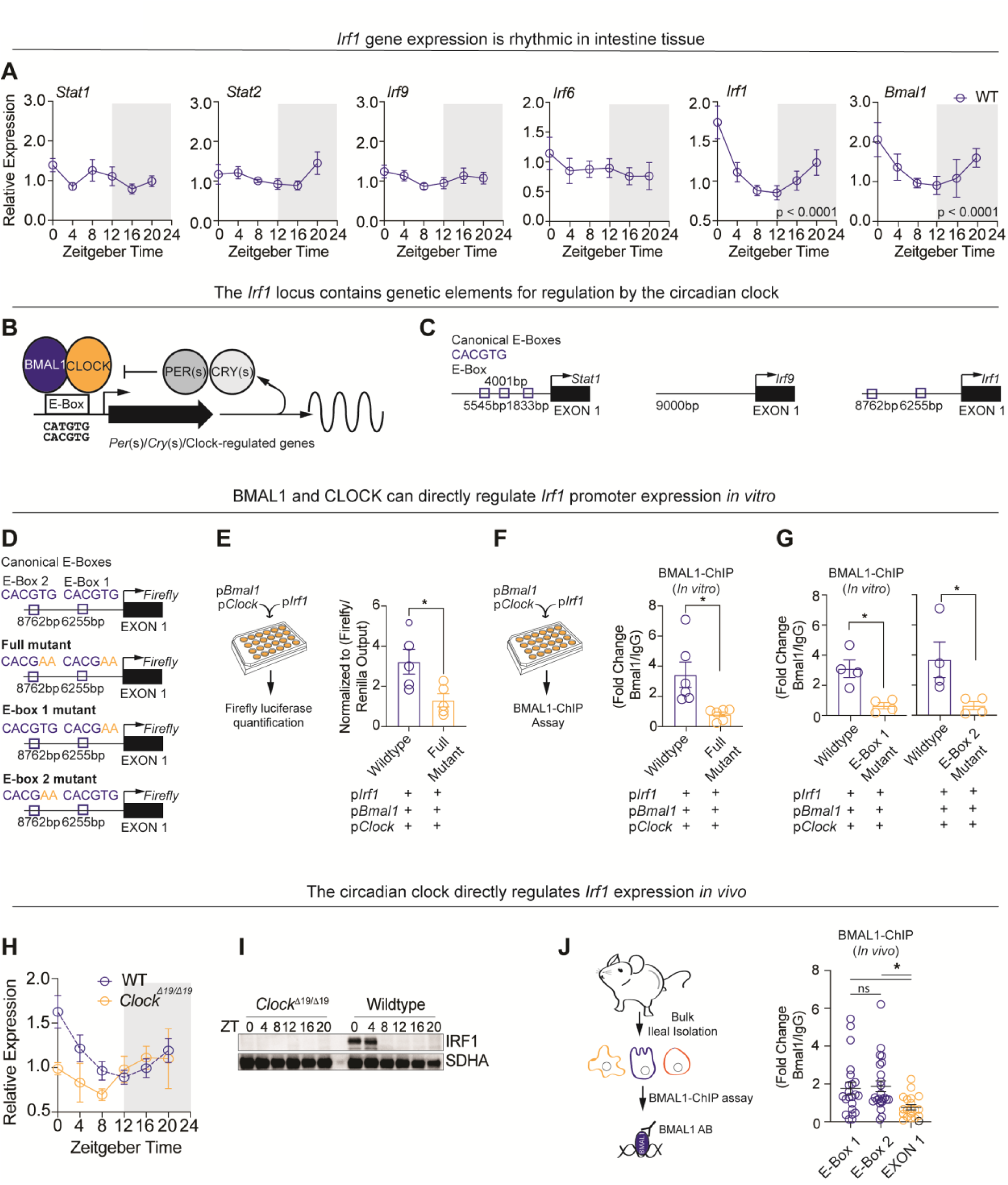
The circadian clock establishes rhythmic expression of the antiviral regulator IRF1. **(A)** qRT-PCR quantification of *Stat1, Stat2,* and select IRFs at six timepoints across the day-night cycle. *Irf1* and *Bmal1* are significantly rhythmic. **(B)** BMAL1 and CLOCK bind E-box elements in promoters to promote transcription of circadian-controlled genes, including their own repressor proteins, PERs and CRYs. **(C)** Putative E-box elements upstream of *Stat1, Irf9,* and *Irf1*. **(D)** Visualization of the *Irf1* promoter showing wildtype E-Boxes, and mutated (Δ) E-Boxes in luciferase reporters. **(E)** Left: Transfection-based luciferase reporter assay. Right: Quantification of *Irf1* promoter activity using a luciferase reporter assay for wildtype vs. mutated E-box elements. **(F)** Left: Transfection-based BMAL1 ChIP assay. Right: Quantification of BMAL1 binding to the *Irf1* promoter *in vitro* for wildtype vs. dual mutated E-box elements. **(G)** Quantification of BMAL1 binding to the *Irf1* promoter *in vitro* for wildtype vs. singly mutated E-box elements (E-box 1 vs. E-box 2). **(H)** qRT-PCR quantification of *Irf1* in wildtype mice versus *Clock ^Δ19/Δ19^* mice at six timepoints across the day-night cycle. **(I)** Western blot examination of IRF1in wildtype mice versus *Clock ^Δ19/Δ19^* mice. **(J)** Quantification of BMAL1 binding to the *Irf1* locus *in vivo.* Small intestine ileal tissue was harvested from wildtype mice at ZT0, followed by BMAL1-ChIP assay targeting E-box 1, E-Box 2 or a downstream control region, EXON1. Data are presented as the ratio of BMAL1 to control IgG antibody. *Stat*, signal transducer and activator of transcription; *Irf*, interferon regulatory factor; BMAL1, brain and muscle ARNT-like 1; *pIrf1, Irf1* plasmid (cloned); *pBmal1, Bmal1* plasmid (constitutively expressed); *pClock, Clock plasmid* (constitutively expressed). Means ± SEM are plotted; *P < 0.05, **P < 0.01; ***P < 0.001; ****P < 0.0001 as determined by Student’s t-test or ANOVA. ns, not significant.

Next, we examined whether *Irf1* expression was directly regulated by circadian clock transcription factors. The core circadian clock operates through a transcriptional-translational feedback loop in which CLOCK and Brain and Muscle ARNT-Like 1 (BMAL1) transcription factors form a heterodimer that binds enhancer (E)-box motifs to regulate gene expression (Fig. 2B). Promoter analysis revealed that *Irf1* contains two conserved E-box elements proximal to its transcriptional start site (Fig. 2C). To test whether *Irf1* is a direct clock-controlled gene, we used luciferase reporter assays and chromatin immunoprecipitation (ChIP) *in vitro* and expression analysis plus ChIP *in vivo*.

For *in vitro* assays, we cloned the *Irf1* promoter upstream of a luciferase reporter and mutated one or both putative E-boxes (Fig. 2D). First, we assessed transcriptional activation in the presence of CLOCK and BMAL1 using luciferase reporter assays in transfected HEK293 cells (Fig. 2E). CLOCK and BMAL1 robustly activated the wildtype *Irf1* promoter, whereas mutation of the E-box elements abolished this effect (Fig. 2E). Next, we assessed binding of BMAL1 to the *Irf1* promoter using ChIP assays. We transfected HEK293 cells with *Irf1* promoter plasmids and CLOCK and BMAL1 expression plasmids, isolated chromatin, immunoprecipitated with anti-BMAL1 or control antibody, and quantified bound DNA using qPCR for the *Irf1* promoter region (Fig. 2F). We found BMAL1-specific binding to the wildtype *Irf1* promoter, but reduced binding to *Irf1* promoter with mutated E-boxes (Fig. 2F-G).

For *in vivo* assays, we examined *Irf1* expression in ileal tissues, and we performed ChIP to assess BMAL1 binding to the *Irf1* promoter. As measured by qRT-PCR, *Irf1* RNA was oscillatory over time in wildtype mice but not in *Clock^Δ19/Δ19^*mice (Fig. 2H). At the protein level, IRF1 levels were high in wildtype mice at ZT0/4 but low at other time points in wildtype mice and at all time points in *Clock^Δ19/Δ19^* mice (Fig. 2I). Furthermore, in *Clock^Δ19/Δ19^*mice, *Irf1* expression was arrhythmic (Fig. S2C). ChIP assays performed with intestinal tissue further demonstrated BMAL1 binding to both E-boxes at the endogenous *Irf1* locus *in vivo,* which was significantly higher than binding to control downstream sequence (EXON1)(Fig. 2J). These data establish *Irf1* as a direct transcriptional target of the circadian clock proteins CLOCK and BMAL1.

### IRF1 is required for temporal gating of antiviral gene expression

To determine whether IRF1 is required for time-of-day–dependent resistance to infection, we examined *Irf1^-/-^* mice (Fig. 3A, Fig. S3A-B). Loss of IRF1 abolished temporal differences in CVB3 replication, with viral titers remaining comparable regardless of infection time (Fig. 3B). Given that food intake influences oscillatory gene expression and knockout mice may have altered feeding behaviors, we quantified food intake in wildtype vs. *Irf1^-/-^*mice. *Irf1^-/-^* mice retained normal feeding rhythms (Fig. 3C, Fig. S3C) and intact oscillations of core clock genes (Fig. 3D, Fig. S3D), indicating that loss of temporal gating did not reflect disruption of circadian behavior or clock function.

**Fig. 3.**
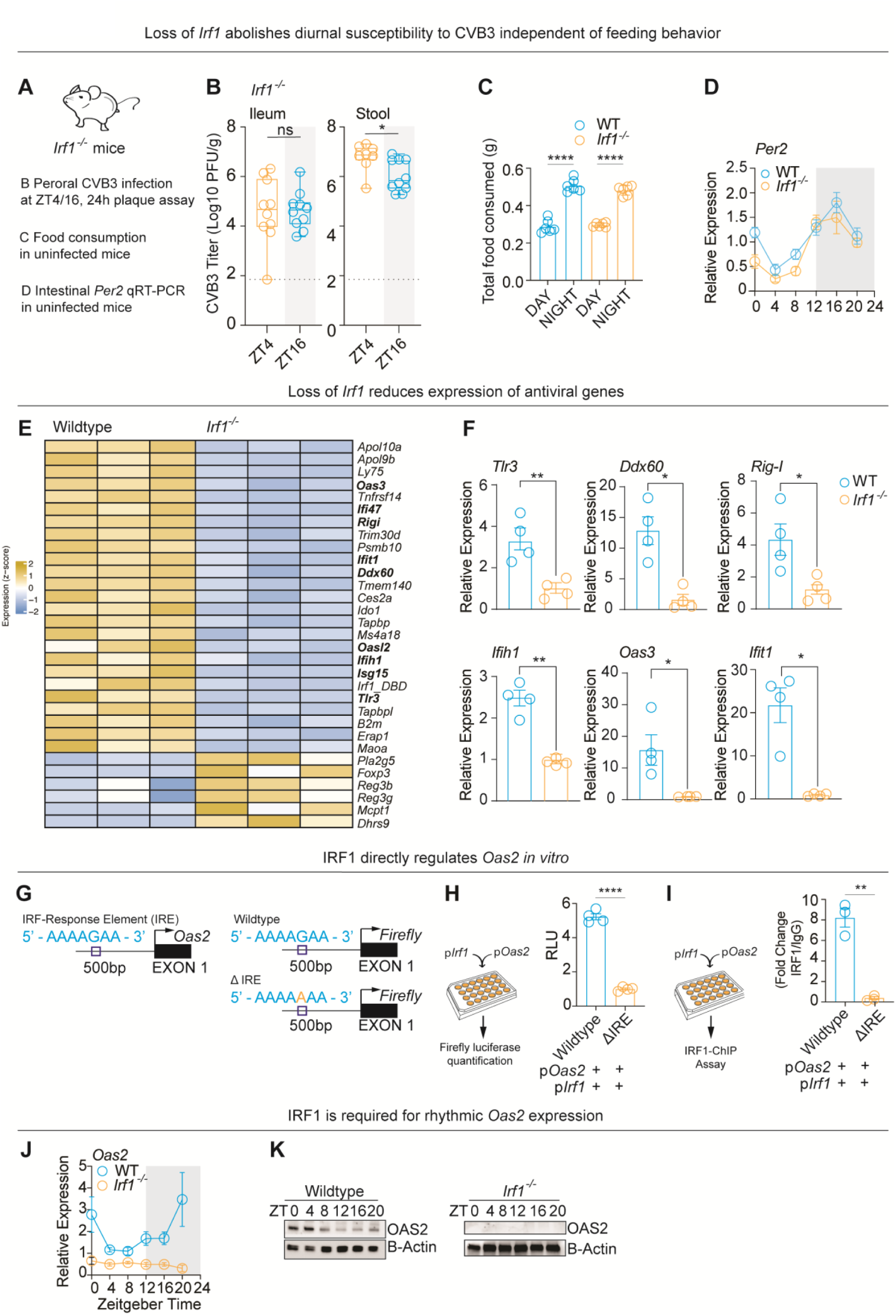
IRF1 is required for temporal gating of antiviral immunity. **(A)** Characterization of *Irf1^-/-^* mice, including peroral infection and titer assay, food consumption quantification, and assessment of circadian gene expression. **(B)** *Irf1^-/-^* mice were perorally infected with 10^9^ PFU CVB3 at one timepoint during the day (ZT4) and one timepoint during the evening (ZT16) and tissues were harvested at 24h post-infection followed by plaque assay using HeLa cells. **(C)** Feeding rhythms were examined in wildtype vs. *Irf1^-/-^* mice across several days. The total amount of food consumed per day and night was quantified. **(D)** qRT-PCR quantification of *Per2* at six timepoints across the day-night cycle in wildtype vs. *Irf1^-/-^* mice. **(E)** Heatmap comparing gene expression in wildtype vs. *Irf1^-/-^* mice from RNA-seq of ZT0 ileum tissue. **(F)** ZT0 qRT-PCR quantification of *Tlr3*, *Ddx60*, *Rig-I*, *Ifih1*, *Oas3*, and *Ifit1* in wildtype versus *Irf1^-/-^* mice. **(G)** Visualization of the *Oas2* promoter showing wildtype IRE, and mutated (Δ) IRE in luciferase reporters. **(H)** Left: Transfection-based *Oas2* luciferase reporter assay. Right: Quantification of *Oas2* promoter activity using a luciferase reporter assay for wildtype vs. mutated IRE. **(I)** Left: Transfection-based IRF1 ChIP assay. Right: Quantification of IRF1 binding to the *Oas2* promoter *in vitro* for wildtype vs. mutated IRE. **(J)** qRT-PCR quantification of *Oas2* at six timepoints across the day-night cycle in wildtype and *Irf1^-/-^* mice. **(K)** Western blot examination of OAS2 at six timepoints across the day-night cycle in wildtype and *Irf1^-/-^* mice. *Per2, Period 2*. *Tlr3*, toll-like receptor 3; *Ddx60*, DExD/H-box helicase 60; *Rig-I*, retinoic acid-inducible gene I; *Ifih1*, interferon induced with helicase C domain 1; *Oas3*, 2’-5’-oligoadenylate synthetase 3; *Ifit1* interferon-induced protein with tetratricopeptide repeats 1; IRE, IRF response element; *Oas2* 2’-5’-oligoadenylate synthetase 2; *pOas2, Oas2* plasmid (cloned); *pIrf1, Irf1* plasmid (constitutively expressed). Means ± SEM are plotted; *P < 0.05, **P < 0.01; ***P < 0.001; ****P < 0.0001 as determined by Student’s t-test or ANOVA. ns, not significant.

Next, we evaluated the antiviral programs downstream of IRF1. First, we performed RNA sequencing from ileal tissue at ZT0, corresponding to peak IRF1 expression in wildtype animals. IRF1-deficiency resulted in reduced expression of multiple genes involved in viral sensing and restriction, including *Oas2*, *Oas3*, *Oasl2*, *Ddx60, Ddx58* (RIG-I), *Ifih1* (MDA5), *Ifit1*, and *Tlr3* (Fig. 3E), indicating that IRF1 sustains a basal antiviral transcriptional state during periods of maximal resistance. Using qRT-PCR, we confirmed decreased expression of these and other immune genes when comparing *Irf1*-deficient mice to wildtype mice (Fig. 3F).

IRF1 binds to IRF-Response Elements (IREs) in promoters of target genes and we used *Oas2* as a representative IRF1 target gene to gain more insight into rhythmic control of innate immune responses. We generated luciferase reporters with wildtype or mutated IREs in the *Oas2* promoter and used them for *in vitro* assays (Fig. 3G-I). IRF1 induced reporter activity from the wildtype promoter, but not from an IRE mutant (Fig. 3H). Next, we assessed binding of IRF1 to the *Oas2* promoter using ChIP assays in HEK293 cells (Fig. 3I). We found IRF1-specific binding to the wildtype *Oas2* promoter, but reduced binding to *Oas2* promoter with a mutated IRE (Fig. 3I). Finally, we quantified expression of *Oas2* in wildtype vs. *Irf1^-/-^* mice. *Oas2* gene and protein expression was minimal in *Irf1^-/-^* mice, but detectable and rhythmic in wildtype mice (Fig. 3J, K, Fig. S3F). Together, these data indicate that IRF1 is required to establish a temporally regulated antiviral state that restricts viral replication.

### A myeloid cell-intrinsic program controls temporal antiviral resistance

We next sought to identify the cellular basis of rhythmic IRF1-dependent antiviral effects. The intestine contains a variety of cell types, including epithelial cells that overlay a large number of immune and other cell types within the lamina propria (Fig. 4A). We first compared *Irf1* expression across intestinal compartments by isolating lamina propria cells vs. epithelial cells from uninfected wildtype mice at ZT0. *Irf1* expression was markedly enriched in lamina propria cells relative to epithelial cells (Fig. 4B). Myeloid cells are abundant (*29*) within the lamina propria and previous work established that *Irf1* is expressed in myeloid populations (*30*). Consistent with this, antibody-mediated depletion of CSF1R-positive myeloid cells at ZT4 significantly reduced *Irf1* expression within the lamina propria (Fig. 4C), identifying myeloid cells as a source of IRF1 in the intestinal immune compartment. Finally, we examined viral loads in mice with lineage-specific deletion of *Irf1* in myeloid cells (*31*), both to narrow the relevant cell types subject to these anticipatory antiviral pathways and to rule out confounding effects of global *Irf1* deletion including impacts on T cell development (*27*). Myeloid-specific loss of IRF1 in *Irf1^ΔLysM^* mice, but not in *Irf1^fl/fl^* controls, abolished time-of-day–dependent differences in CVB3 titers, phenocopying global *Irf1* deficiency (Fig. 4D). Together, these data demonstrate that myeloid cell-intrinsic IRF1 expression, driven by circadian transcription factors, establishes daily rhythms in antiviral programs.

**Fig. 4.**
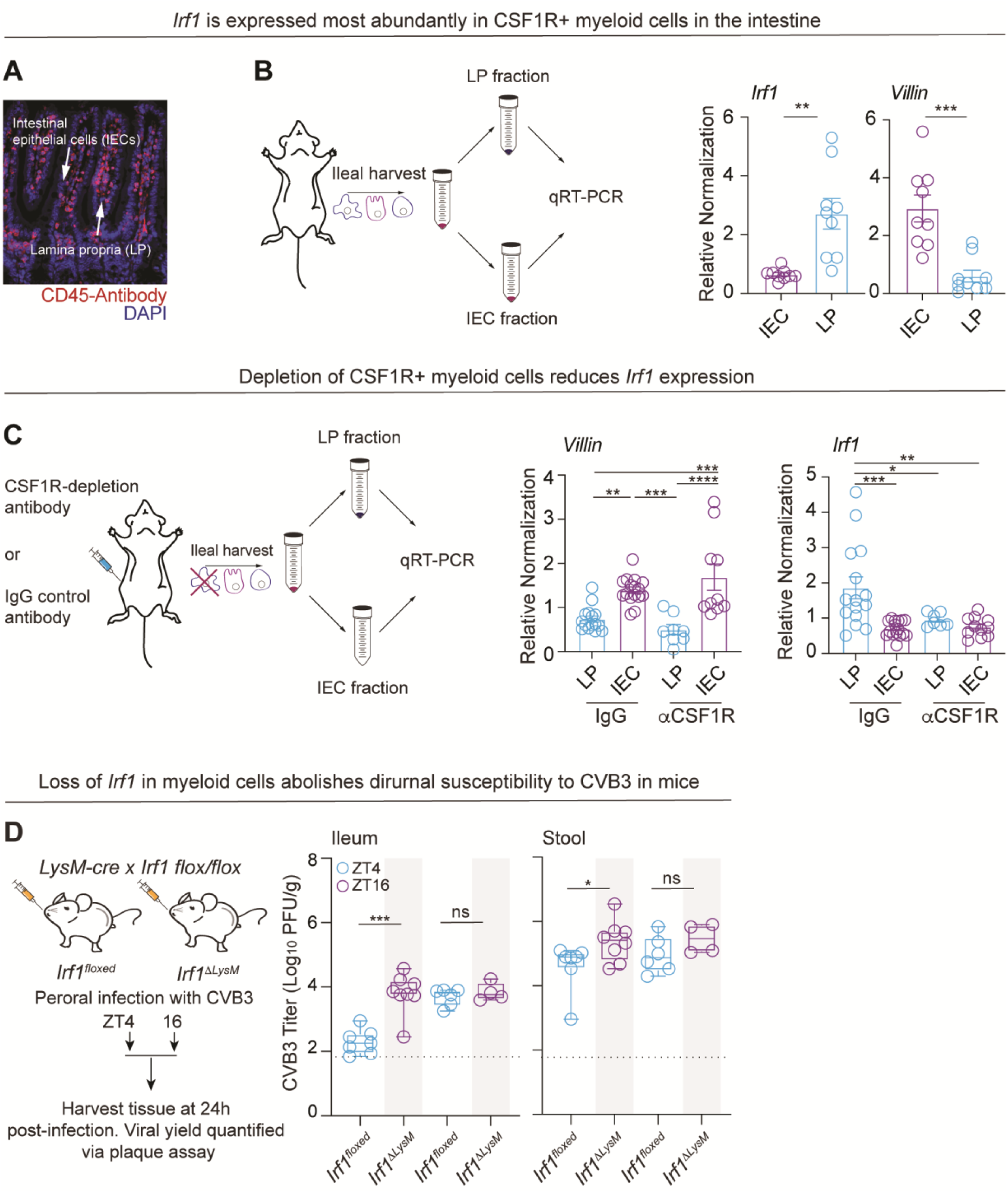
A myeloid cell–intrinsic program controls temporal antiviral resistance. **(A)** Microscopy of murine small intestine, showing intestinal epithelial cells (IECs) and lamina propria (LP) cells (stained with CD45 antibody) beneath. **(B)** Gene expression in IEC vs. LP. Small intestine from wildtype was collected at ZT4, IECs and LP cells were purified, and qRT-PCR quantified expression of *Irf1* and *Villin*, a marker of IECs. **(C)** Gene expression in mice with depleted CSF1R+ cells. Wildtype mice were injected at ZT4 with a macrophage depletion antibody (anti-CSF1R) or IgG control and 72h later intestine was collected, IECs and LP cells were isolated, and qRT-PCR quantified expression of *Irf1* and *Villin*. **(D)** Viral load in the ileum and stool of perorally infected mice deficient for *Irf1* specifically in myeloid cells (*Irf1^ΔLysM^*) and controls (*Irf1^floxed^*). Mice were infected with 10^9^ PFU CVB3 at one timepoint during the day (ZT0) and one timepoint during the evening (ZT12) followed by quantification of viral load by plaque assay. IECs, intestinal epithelial cells; LP, lamina propria; CSF1R Colony-stimulating factor 1 receptor; IgG, immunoglobulin G. Means ± SEM are plotted; *P < 0.05, **P < 0.01; ***P < 0.001; ****P < 0.0001 as determined by Student’s t-test or ANOVA. ns, not significant.

## DISCUSSION

Our findings establish that host susceptibility to enteric viral infection oscillates by time of day through anticipatory antiviral responses. We identify a myeloid cell–intrinsic program that controls expression of the antiviral transcription factor IRF1, establishing daily rhythms in basal antiviral gene expression and defining temporal windows of host resistance. These results reveal time of infection as an underappreciated determinant of viral outcome, comparable to host genetics, immune status, or route of exposure, and highlight the importance of reporting time of day for reproducibility of infection studies. Notably, anticipatory immune regulation has also been described in plants, where circadian programs precondition defense pathways prior to pathogen exposure (*32*). Together, these findings suggest that anticipatory immunity may represent a conserved, cross-kingdom strategy aligning host defense with predictable environmental risk.

Our data suggest that clock control acts early in infection. Viral titers diverge within the first 24 hours, consistent with effects on the initial cycles of replication. Rhythmic IRF1 and downstream antiviral gene expression occur prior to infection, supporting a basal antiviral state rather than an inducible response. The IRF1-dependent gene set includes sensors and restriction factors such as MDA5(*33*), OAS family members (*34*) and TLR3 (*35, 36*), indicating coordinated regulation of early antiviral defenses. Notably, time-of-day differences are observed following oral but not systemic infection, indicating that temporal control operates at the mucosal interface during infection by the natural oral route. Together, these findings support a model in which timing shapes the earliest host–virus interactions, biasing infection outcome before widespread replication is established. This mode of regulation contrasts with prior clock-virus studies, which have largely focused on how time of day influences host responses after infection or effects on viral replication dynamics (*12–16*). Here, we show that the circadian clock instead programs antiviral immunity prior to infection, establishing a basal state that gates susceptibility at the moment of exposure.

Clock regulation of antiviral immunity operates through a cell-type–specific transcriptional circuit that establishes a transient antiviral state. We identify CSF1R-positive myeloid cells in the intestinal lamina propria as a major source of rhythmic IRF1 expression, positioning these cells as temporal sentinels that preemptively shape the antiviral landscape. Within this compartment, the circadian clock selectively targets IRF1 rather than broadly oscillating interferon pathways, focusing regulation on a single transcriptional node that sustains basal antiviral gene expression. This architecture may balance immune readiness with the metabolic and inflammatory costs of sustained activation, enabling anticipatory protection without continuous immune stimulation (*37*).

The emergence of a temporally gated basal antiviral state in mice raises the question of why such anticipatory immunity arose and is maintained. One possibility is that this program reflects evolutionary selection imposed by recurrent or predictable pathogen exposure at specific times of day (*37*). In this model, individuals with elevated antiviral defenses aligned with periods of increased exposure risk would have enhanced survival, leading to retention of anticipatory immunity. Limiting immune activation to defined temporal windows may optimize the tradeoff between protection and cost, minimizing metabolic burden and immunopathology while preserving effective host defense. In nocturnal animals such as mice, this anticipatory immune response may be coordinated with behavior and infection dynamics. Following oral exposure, likely during the active dark phase, productive viral replication in the intestine is delayed by several hours due to gastrointestinal transit (*21*), creating a temporal separation between exposure and replication. Subsequently, in the early light phase, peak antiviral gene expression aligns with inefficient viral replication. This temporal offset positions viral replication within a pre-existing antiviral state, allowing a clock driven antiviral program to restrict early viral replication.

In summary, our work reveals a rhythmic innate immune program that establishes antiviral resistance through IRF1, redefining host susceptibility as a dynamic and anticipatory state. This framework may help explain variability in infection outcomes not accounted for by exposure, genetics, or immunocompetence, and has implications for understanding viral transmission and optimizing the timing of interventions.

## Acknowledgments

We thank Dr. Lora Hooper and Dr. Joseph Takahashi for their feedback, Dr. Carla Green for access to feeding cages, Laboratory Animal Resources (Princeton) for mouse maintenance, and the Genomics Core Facility (Princeton) for their support with sequencing.

## Author contributions

Conceptualization: TOO, RWM, JKP, JFB

Methodology: TOO, RWM, MWA, BTM, HD, IJ, DAS, UB, CD, VLT, JKP, JFB

Investigation: TOO, RWM, MWA, BTM, HD, IJ, DAS, UB, CD, VLT, JKP, JFB

Visualization: TOO, RWM, JKP, JFB

Funding acquisition: JKP, JFB

Project administration: JKP, JFB

Supervision: JFB, JKP

Writing – original draft: TOO, RM, JP, JFB

Writing – review & editing: TOO, RM, MWA, BM, HD, IJ, DAS, UB, CD, VLT, JP, JFB

## Data, code, and materials availability

Bulk RNA-sequencing data is available here: https://osf.io/te8am/overview?view_only=8193460b964c4379b679bc2d29c53952

## Supplementary Materials

### Materials and Methods

#### Virus and cells

HeLa cells were grown at 37°C in Dulbecco’s Modified Eagle Medium (DMEM) media supplemented with 10% newborn calf serum (Fisher Scientific, #NC1928575) and Penicillin-Streptomycin (10,000U penicillin, 10mg streptomycin, Millipore Sigma, #P4333-100ml). A CVB3-H3 stock (a gift from Marco Vignuzzi; GenBank: U57056.1) was generated by co-transfecting an infectious clone plasmid with a T7 RNA polymerase expression plasmid into HeLa cells. Further amplification was performed to generate a high-titer viral stock. Plaque Forming Units (PFU) were quantified by plaque assay using HeLa cells (*38*).

#### Mice

All animal procedures followed NIH guidelines and were approved by the University of Texas Southwestern Medical Center IACUC (Animal Welfare Assurance A3472-01) or Princeton University IACUC (Animal Welfare Assurance D16-00273). All work was performed with the design in mind for minimal pain and animal welfare at the forefront. In the event of severe disease or distress, mice were promptly euthanized. C57BL/6J mice and mice *Irf1^-/-^* mice were obtained from Jackson Laboratory (Strain: #000664 and #002762, respectively). Additionally, *Irf1*^flox/flox^ mice (V. Tarakanova, Medical College of Wisconsin) mice were crossed with LysM Cre mice (Strain: #004781) to generate *Irf1^ΔLysM^*and *Irf1^fl/fl^* littermate controls for experimental use. Whole body non-functional *Clock*^Δ19^ mice (missing exon 19 in CLOCK; dominant negative mutation) were obtained from J. Takahashi (UT Southwestern Medical Center). Mice were housed in 12-hour light/ 12-hour dark cycle within specific pathogen-free (SPF) barrier facilities prior to experiments. 8–12-week-old mice were used in all experiments. Any modulations of lighting conditions used isolation cabinets with green LED lights programmed by ClockLab v3.604 and Chamber Control Software v4.114 (Actimetrics Inc., Wilmette, IL).

#### Infections and plaque assays

Mice were infected by peroral infection (10^9^ plaque forming units of CVB3-H3 in 25 ul pipetted into the mouth) or intraperitoneal injection (10^5^ plaque forming units of CVB3-H3 in 100 ul). Additionally, due to a previously published a sex bias in murine CVB3 infection, only male mice were used for infection experiments (*26*). All infections took place at Zeitgeber times (ZT) 0, 4, 8, 12, 16, or 20. 24 hours after infection, ileum, cecum, and stool samples from each mouse were harvested for further analysis. At 24h post-infection, tissues or stool were weighed and homogenized using a bead beater homogenizer, followed by three freeze-thaw cycles to release virus prior to plaque assay using HeLa cells (*39*). Briefly, dilutions of virus were used to infect HeLa cell monolayers which were then covered with 1% agarose overlays. At two days post infection, cells were fixed and stained with a crystal violet solution and plaques were counted.

#### qRT-PCR Gene Expression Experiments

Murine samples were harvested and placed in 1mL TRIzol™ Reagent. To extract RNA, samples were homogenized and extracted using the Qiagen RNeasy^®^ Plus Universal Mini Kit. cDNA synthesis began by adding 10ng of sample RNA to water (for 14 μL total), 1 μL Random Hexamers, 1 μL dNTP, and heating to 65°C for 5 minutes. Samples were then immediately chilled on ice for 2 minutes. Next, to each tube 5 μL 5x first-stand buffer, 2 μL 0.1M DTT, and 1 μL RNAaseOUT were added to each sample and incubated at 37°C for 2 minutes. Following this, 1 μL of M-MLV RT was added to each sample and the tubes were incubated using the MMV1 method. Following cDNA synthesis, samples were examined with quantitative PCR. For each reaction, 1 μL of cDNA was added to 10 μL Platinum™ SYBR™ Green qPCR SuperMix-UDG, 1 μL BSA (20x 1mg/mL), 5.9 μL nuclease free water, 0.1μL ROX Reference Dye, 1 μL of 0.5 μM forward primer, and 1 μL of 0.5 μM reverse primer. Each 20 μL reaction was added into a 384 well plate and incubated using the QS7Pro-384-Well-PCR-Melt-Std method on the Quant Studio™ 7 Pro qPCR machine. Cq scores were analyzed using Design & Analysis (v1 4.3 download) as well as Microsoft Excel (Version 2510, build 19328.20266).

#### Cloning

gBlock sequences were obtained from IDT. Sequences were cloned into a luciferase reporter vector using the New England Biolabs Gibson Assembly® Master Mix. Following colony qPCR and plasmid purification via the Qiagen QIAprep® Spin Miniprep Kit, plasmids from candidate colonies were sent for Sanger sequencing to GENEWIZ from Azenca. Candidate plasmids were aligned to desired sequence and matching sequences were preserved as glycerol stocks.

#### Luciferase Reporter Assays

Human Embryonic Kidney (HEK) cells were plated on a 24-well dish and grown to 70% confluency. Cells were transfected with 500 ng of each plasmid using the FuGENE® HD Transfection Reagent. A Human Luciferase/Renilla Luciferase plasmid and untransfected cells were used as controls. 48 hours after the initial transfection, Firefly and Renilla Luciferase luminescence was measured in each well using the Dual-Glo® Luciferase Assay System, a Tecan Infinite® 200 PRO plate reader, and the i-control™ software.

#### *In vitro* chromatin immunoprecipitation (ChIP)

Human Embryonic Kidney (HEK) cells were plated on a 6-well dish and grown to 70% confluency. Cells were transfected with 1,000ng of each plasmid using the FuGENE® HD Transfection Reagent. 48 hours after the initial transfection, cells were fixed in 1% paraformaldehyde at 37°C for 10 minutes, quenched in 1M glycine at room temperature for 10 minutes, and then washed in PBS+ 1x protease inhibitor + 1x phosphatase inhibitor. Next, using an Eppendorf rotator, cells were lysed in Lysis Buffer 1 (4°C for 10 minutes, then centrifuged at 4°C, 1,350g for 5 minutes) and Lysis Buffer 2 (room temperature for 10 minutes, centrifuged at 4°C, 1,350g for 5 minutes). Following this, cells were resuspended in Lysis Buffer 3 (ThermoFisher Scientific, #00-4333-57), sonicated (Diagenode Bioruptor Plus) for 9 cycles at 20 seconds on and 30 seconds off, and centrifuged at 4°C, 1,350g for 5 minutes. To solubilize the nuclear membrane, 1/10 volume of 10% Triton X-100 was then added to samples. To crack the nuclei, samples were centrifuged at 4°C, 20,000g for 10 minutes and the supernatant (nuclear content) was retained. DNA concentration was quantified using the Qubit 1X dsDNA kit. The chromatin immunoprecipitation reaction consisted of 1 mg of shared chromatin and 2.5 mg of either BMAL1 (Cell Signaling Technology, D2L7G Rabbit Monoclonal Antibody #14020), or IgG (Cell Signaling Technology, Normal Rabbit IgG #2729). The initial reaction was incubated at room temperature for 1 hour on the Eppendorf rotator. Next, Pierce™ Protein A/G Magnetic Beads were cleared in TBS-T buffer and then added to each chromatin immunoprecipitation reaction (25 μL) for 1 hour. The beads were then washed in RIPA buffer (ThermoFisher Scientific, #89901), washed in 1 mL TE 50 mM NaCl, and then eluted in Pierce™ Gentle Ag/Ab Elution Buffer, pH 6.6 at 65°C for 30 minutes. To reverse crosslinks, samples were incubated in RNase A at 37°C for 30 minutes and then incubated in Proteinase K at 57°C overnight. The following day, the DNA was purified using the Qiagen QIAquick^®^ PCR Purification Kit and analyzed by qPCR primer probes (for CLOCK/BMAL1 E-boxes or IRF Response Element). Cq scores were analyzed using Design & Analysis (v1 4.3 download) as well as Microsoft Excel (Version 2510, build 19328.20266), and analyzed using the fold-change method.

#### *In vivo* chromatin immunoprecipitation (ChIP)

Murine intestinal samples were harvested, cut into 0.5cmx0.5cm sections, and washed in cold PBS. Next, samples were placed in 15mL conical tubes, fixed in 1% paraformaldehyde at 37°C for 10 minutes, quenched in 1M glycine, pH 3.2, at room temperature for 10 minutes, and then washed in PBS+ 1x protease inhibitor + 1x phosphatase inhibitor. Next, using an Eppendorf rotator, samples were lysed in Lysis Buffer 1 [consisting of: 50 mM HEPES-KOH (pH 7.5; HEPES: ThermoFisher Scientific, Gibco™, 15630080; KOH: ThermoFisher Scientific, 437135000), 140 mM NaCl (ThermoFisher Scientific, AAJ2161836), 1 mM EDTA (ThermoFisher Scientific, Invitrogen™, 15575020), 10% glycerol (ThermoFisher Scientific, A16205.0F), 0.5% NP-40 (ThermoFisher Scientific, 85124), and 0.25% Triton X-100 (ThermoFisher Scientific, 85111), prepared in ddH₂O.] (4°C for 10 minutes, then centrifuged at 4°C, 1,350g for 5 minutes) and Lysis Buffer 2 [consisting of 10 mM Tris-HCl (pH 8.0; ThermoFisher Scientific, Invitrogen™, 12090015), 200 mM NaCl (ThermoFisher Scientific, AAJ2161836), 1 mM EDTA (ThermoFisher Scientific, Invitrogen™, 15575020), and 0.5 mM EGTA (Millipore Sigma, 324626-25GM), supplemented with protease and phosphatase inhibitors (Halt™ Protease Inhibitor Cocktail, 100X, 78430; Halt™ Phosphatase Inhibitor Cocktail, 78420; ThermoFisher Scientific), prepared in ddH₂O] (room temperature for 10 minutes, centrifuged at 4°C, 1,350g for 5 minutes). Following this, cells were resuspended in Lysis Buffer 3, sonicated (Diagenode Bioruptor Plus) for 12 cycles at 30 seconds on and 30 seconds off, and centrifuged at 4°C, 1,350g for 5 minutes. To solubilize the nuclear membrane, 1/10 volume of 10% Triton X-100 was then added to samples. To crack the nuclei, samples were centrifuged at 4°C, 20,000g and the supernatant (nuclear content) was retained. DNA concentration was quantified using the Qubit 1X dsDNA kit. The chromatin immunoprecipitation reaction consisted of 1 mg of shared chromatin and 2.5 mg of either BMAL1 (Cell Signaling Technology, D2L7G Rabbit Monoclonal Antibody #14020), or IgG (Cell Signaling Technology, Normal Rabbit IgG #2729). The initial reaction was incubated at room temperature for 1 hour on the Eppendorf rotator. Next, Pierce™ Protein A/G Magnetic Beads were cleared in TBS-T buffer and then added to each chromatin immunoprecipitation reaction (25 μL) for 1 hour. The beads were then washed in RIPA buffer, washed in 1 mL TE 50 mM NaCl, and then eluted in Pierce™ Gentle Ag/Ab Elution Buffer, pH 6.6 at 65°C for 30 minutes. To reverse crosslinks, samples were incubated in RNase A at 37°C for 30 minutes and then incubated in Proteinase K at 57°C overnight. The following day, the DNA was purified using the Qiagen QIAquick^®^ PCR Purification Kit and analyzed by qPCR primer probes (for *Irf1* E-boxes and Exon 1). Cq scores were analyzed using Design & Analysis (v1 4.3 download) as well as Microsoft Excel (Version 2510, build 19328.20266), and analyzed using both the fold-change and percent input methods.

#### Western Blot

Mouse ileal tissue was homogenized in T-PER Tissue Protein Extraction Reagent+ protease + phosphatase inhibitor tablets. For each sample, 40 mg of protein was loaded into a 4-20% Mini-PROTEAN® TGX™ protein gel and transferred to a PVDF membrane. Next, membranes were blocked in 5% dry nonfat milk in PBS-T for 1 hour. Membranes were incubated in the following primary antibodies (Cell Signaling Technology, D5E4 (IRF1) Monoclonal Antibody #8478; Cell Signaling Technology, OAS2 Antibody #54155) at 4°C overnight in 1:1,000 dilutions. The following day, membranes were washed in PBS-T. Membranes were then incubated in HRP-conjugated secondary antibody (Cell Signaling Technology, Anti-rabbit IgG, HRP-linked Antibody #7074) at room temperature for 2 hours in 1:10,000 dilutions. A Bio-Rad ChemiDoc™ Imaging System was used to visualize membranes.

#### Food Consumption Measurements

Food intake and feeding times were measured using a metabolic cage (Labmaster, TSE Systems GmbH, Germany). Wildtype and *Irf1^-^*^/-^ mice were individually housed in a light and temperature (22.5–23.5°C) controlled environment. Following a five-day acclimation in the home cage, mice were analyzed in the metabolic chambers for six days. All mice were provided with food (LabDiet, 5K52) and water ad libitum; food and water intake were continuously recorded using lid-mounted sensors.

#### RNA-seq

Murine ileal samples were harvested at ZT0 and placed in 1mL TRIzol™ Reagent. To extract RNA, samples were homogenized and extracted using the Qiagen RNeasy^®^ Plus Universal Mini Kit. Sample quality was analyzed using the Qubit™ RNA High Sensitivity (HS) and the Agilent 2100 Bioanalyzer. Following quality control, an RNA-seq Directional Library was prepared on the Apollo 324 robot. Next, bulk RNA-sequencing was performed using the NovaSeq SP 100nt Flowcell v 1.5 protocol. Following a FastQC file quality check, sample reads were aligned to the UCSC reference mouse genome (mm39) in R (version 4.5.2). Differential Gene Expression Counts were generated using the DESeq2 package in R. Reads with low mapping quality (less than 10 recounts) were filtered out.

#### Lamina Propria and CSF1R Depletion Experiment

For the macrophage depletion experiment, wildtype mice were injected at ZT4 with 250 μL of antibodies to the CSF1R receptor (BioXCell, BE0213) or IgG as a control (BioXCell, BE0089). 24 hours after the initial injection, all mice were injected again with the same antibody as the previous day. 24 hours after the second injection, animals were sacrificed and terminal ileum samples were collected for a lamina propria isolation.

Murine terminal ileum was collected and placed in PBS+2%FBS; Peyer’s patches were removed from samples. IECs were isolated using EDTA-DTT Buffer at 37°C for 15 minutes. IECs were placed in TRIzol™ Reagent for subsequent gene expression analysis. To isolate the lamina propria, samples (not containing IECs) were incubated in Digest Solution at 37°C, 250 RPM for 1 hour. Using the 40% and 80% Percoll solutions, a Percoll gradient was established to extract cells in the lamina propria. Lamina propria cells were placed in TRIzol™ Reagent for subsequent gene expression analysis.

#### Promoter Analysis

Promoter analysis for E-boxes and IREs was performed using the Genome Reference Consortium Mouse Build 39 (GRCm39/mm39) reference genome in the University of California, Santa Cruz browser.

#### Statistical Analysis

Plaque assays for two conditions were examined for normality to determine which unpaired T-test (Mann-Whitney) was used. Plaque assays using more than two conditions used one-way ANOVA with multiple comparisons with Tukey corrections. Furthermore, CircaCompare was used to detect rhythmicity, determine peak and trough of the data, and establish a best-fit cosinor line values. Rhythms with a p-value less than 0.0001 were defined as significantly rhythmic (*40*).

## Figure Legends

**Fig. S1.**
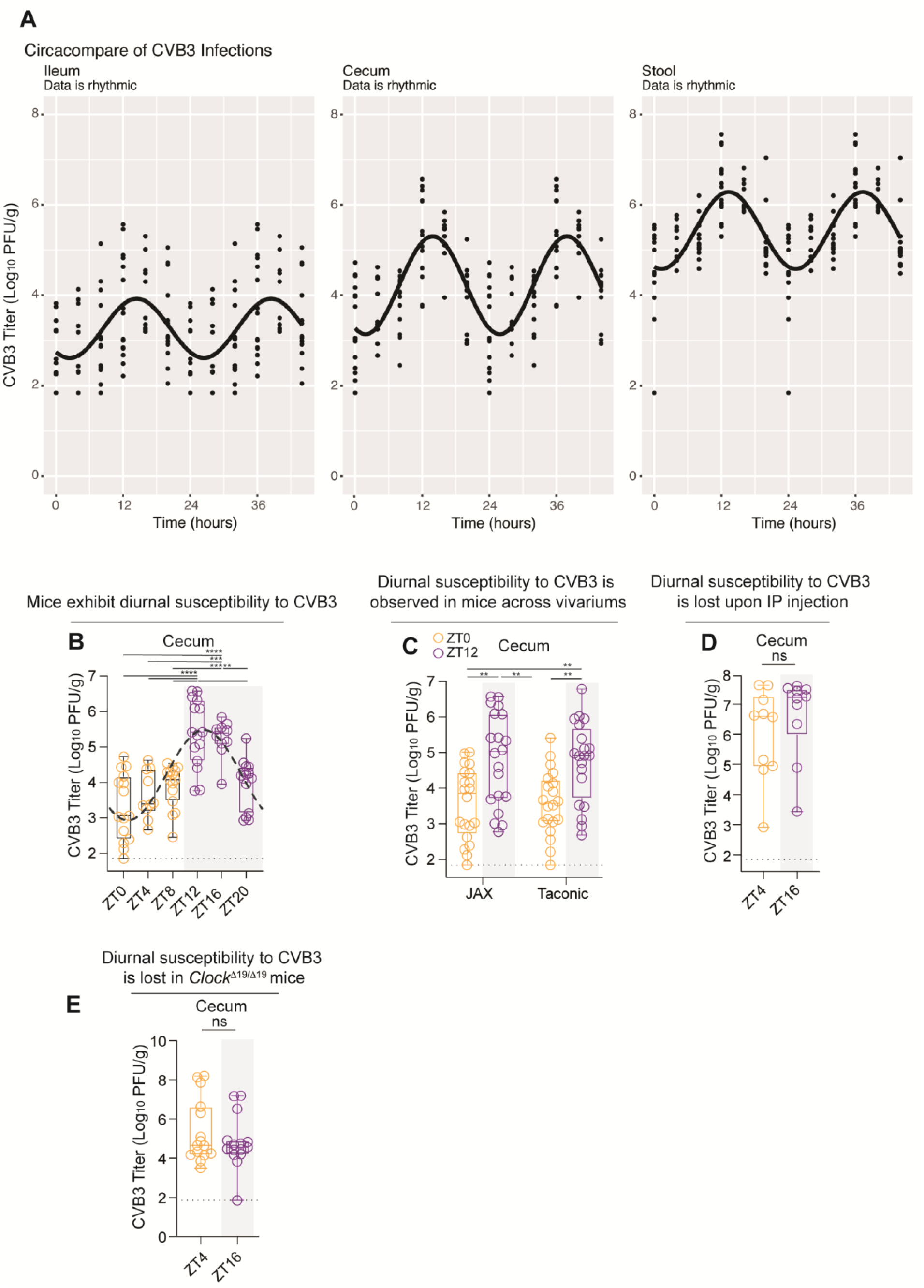
Time of day determines host susceptibility to enteric viral infection in the cecum. (**A**) CircaCompare analysis of ileum, cecum, and stool following peroral infection with 10^9^ PFU CVB3 at six timepoints across the day-night cycle. Data was double plotted for visualization and rhythmicity was assessed via CircaCompare in R (P < 0.0001). **(B)** Mice were perorally infected with 10^9^ PFU CVB3 at six timepoints across the day-night cycle and viral load was quantified in the cecum at 24h post-infection. Black dashed line shows rhythmicity analysis of peak and trough within the data set. Gray dotted line indicates detection limit. **(C)** Quantification of viral load in cecum 24h post-infection in wildtype C57BL/6 mice from The Jackson Laboratory versus Taconic Biosciences. **(D)** Quantification of viral load in the cecum 24h post-infection following intraperitoneal (IP) injection in wildtype mice. **(E)** Quantification of viral load in the cecum 24h post infection in *Clock^Δ19/Δ19^* mice. Best fit curves for rhythmicity are plotted. ****P < 0.0001 as determined by CircaCompare.

**Fig. S2.**
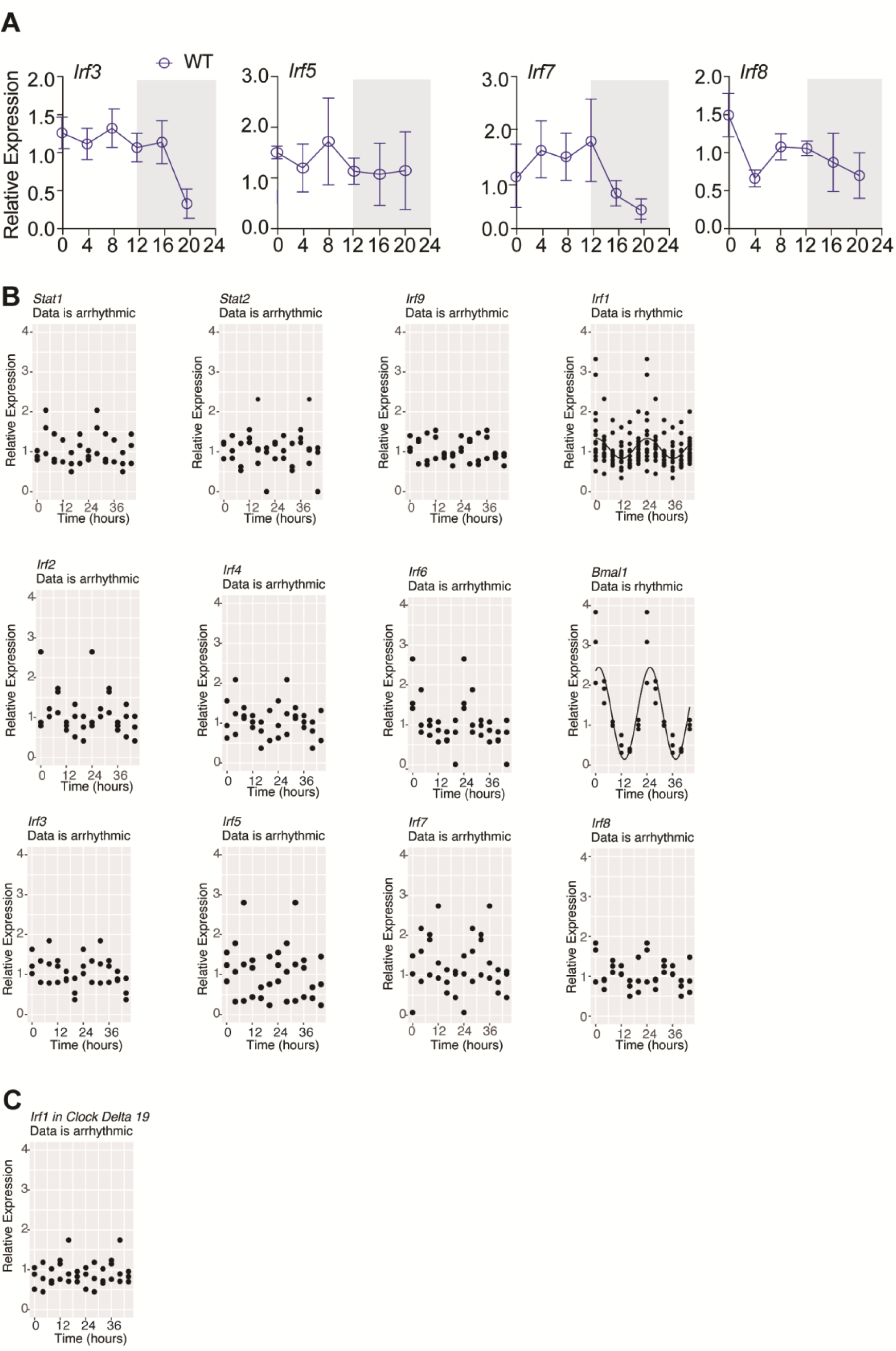
Expression of certain host genes is rhythmic in uninfected mice. (**A**) qRT-PCR quantification of *Irf3, Irf5, Irf7,* and *Irf8* at six timepoints across the day-night cycle. **(B)** CircaCompare analysis of gene expression for *Bmal1* and antiviral genes (*Stat1*, *Stat2*, and *Irf1-Irf9).* Data was double plotted for visualization and rhythmicity was assessed via CircaCompare in R. **(C)** CircaCompare analysis of *Irf1* gene expression in *Clock ^Δ19/Δ19^* mice. Data were double plotted for visualization and rhythmicity was assessed via CircaCompare. Best fit curve for rhythmic genes are plotted. ****P < 0.0001 as determined by CircaCompare.

**Fig. S3.**
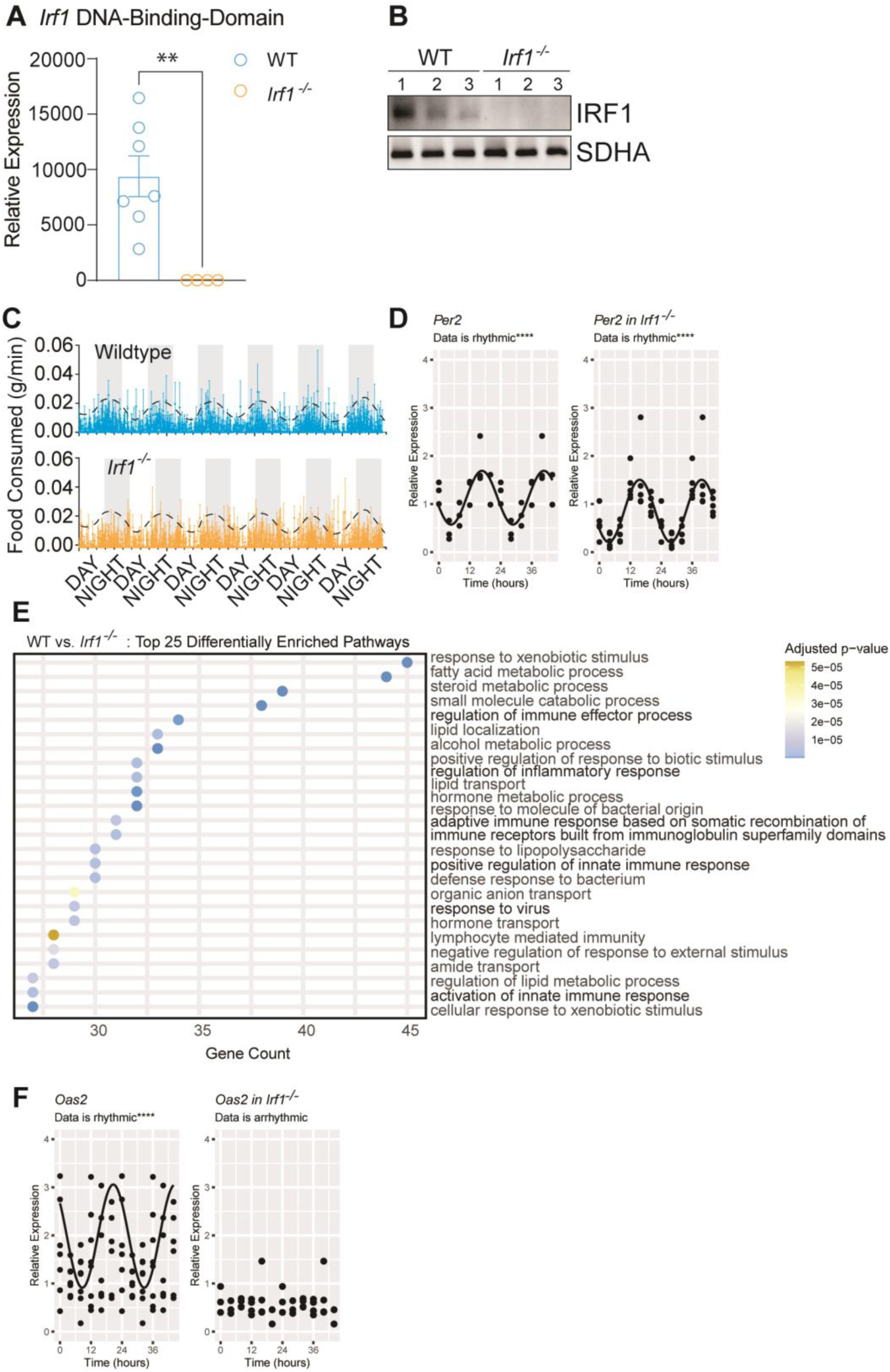
Confirmation of IRF1 loss and circadian clock preservation in *Irf1^-/-^* mice. **(A)** qRT-PCR quantification of the *Irf1* DNA-Binding Domain at ZT0 in wildtype and *Irf1^-/-^* mice. **(B)** Western blot quantification of IRF1 at ZT0 in wildtype and *Irf1^-/-^* mice. **(C)** Food consumed across the day-night cycle in wildtype and *Irf1^-/-^* mice. **(D)** CircaCompare examination of *Per2* across the day-night cycle in wildtype (P < 0.0001) and *Irf1^-/-^* (P < 0.0001) mice. **(E)** Gene ontology for Top 25 differentially enriched pathways in wildtype vs. *Irf1^-/-^* mice. **(F)** CircaCompare examination of *Oas2* across the day-night cycle in wildtype (P < 0.0001) and *Irf1^-/-^* (P < 0.0001) mice. Means ± SEM are plotted; **P < 0.01; as determined by Student’s t-test. For rhythmic data, best fit curves for rhythmicity are plotted. ****P < 0.0001 as determined by CircaCompare.

**Table S1:** Mice.

| Item | Vendor | Catalog Number |
| --- | --- | --- |
| C57BL/6J wildtype Mice | The Jackson Laboratory | 00664 |
| B6.129S2-Irf1<sup>tm1Mak</sup>/J | The Jackson Laboratory | 002762 |
| B6.129-Bmal1<sup>tm1Bra</sup>/J | The Jackson Laboratory | 009100 |
| C57BL/6J Irf1<sup>fl/fl</sup> | Dr. Vera Tarakanova | NA |
| C.B6-Clock<sup>tm1Jt</sup>/J | The Jackson Laboratory | 016175 |
| B6.129P2-Lyz2<sup>tm1(cre)</sup>Ifo/J | The Jackson Laboratory | 004781 |
| C57BL/6NTac | Taconic Biosciences | B6-M |

**Table S2:** Cell Culture and Virus Infection Reagents.

| Item | Vendor | Catalog Number |
| --- | --- | --- |
| HeLa Cells | Dr. Karla Kirkegaard | NA |
| CVB3 H3 Clone | Dr. Marco Vignuzzi | NA |
| Dulbecco's Modified Eagle's Medium (DMEM), High Glucose | ThermoFisher Scientific | 11965118 |
| Newborn Calf Serum (Heat Inactivated) | ThermoFisher Scientific | NC1928575 |
| Lipofectamine 2000 Transfection Reagent 1.5mL | ThermoFisher Scientific | 11668019 |
| Phosphate-buffered saline, no calcium or magnesium (PBS) | ThermoFisher Scientific, Gibco™ | 14190144 |

**Table S3:** Tissue Processing and Plaque Assay Reagents.

| Item | Vendor | Catalog Number |
| --- | --- | --- |
| Fisherbrand™ Bead Mill 4 Mini Homogenizer | Fisherbrand™ | 15-340-164 |
| Omni Bulk Beads, Stainless Steel 2.4mm | Millipore Sigma | MNI-19-640 |
| Dulbecco's phosphate-buffered saline with calcium and magnesium (PBS+) | ThermoFisher Scientific, Gibco™ | 14040133 |
| Phosphate-buffered saline, no calcium or magnesium (PBS) | ThermoFisher Scientific, Gibco™ | 14190144 |
| Dulbecco's Modified Eagle's Medium (DMEM), High Glucose | ThermoFisher Scientific | 11965118 |
| Sodium bicarbonate powder 500g | Millipore Sigma | S8875-500G |
| SeaKem LE Agarose 500g | Lonza | BMA50000 |
| Crystal Violet | Millipore Sigma | C0775-25G |

**Table S4:** qRT-PCR Gene Expression Experiments Reagents.

| **Item** | **Vendor** | **Catalog Number** |
| --- | --- | --- |
| TRIzol™ | ThermoFisher Scientific, Invitrogen™ | 15596026 |
| Qiagen RNeasy® Plus Universal Mini Kit | Qiagen | 73404 |
| Random Hexamers 50 uM | ThermoFisher Scientific, Invitrogen™ | N8080127 |
| 10 mM dNTP Mix | ThermoFisher Scientific, Invitrogen™ | 18427013 |
| 5x First-Strand Buffer | ThermoFisher Scientific, Invitrogen™ | 18057018 |
| DTT | ThermoFisher Scientific, Invitrogen™ | 18057018 |
| M-MLV RT | ThermoFisher Scientific, Invitrogen™ | 18057018 |
| RNaseOUT™ Recombinant Ribonuclease Inhibitor | ThermoFisher Scientific, Invitrogen™ | 10777019 |
| Platinum™ SYBR™ Green qPCR SuperMix-UDG (includes ROX and BSA) | ThermoFisher Scientific, Invitrogen™ | 11733046 |
| ROX Reference Dye | ThermoFisher Scientific, Invitrogen™ | 12223012 |
| HyClone™ HyPure Water, Molecular Biology Grade | Cytiva | SH30538.FS |
| Applied Biosystems™ QuantStudio™ 7 Pro Real-Time PCR System, 384-well | Fisher Scientific | A43171 |

**Table S5:**
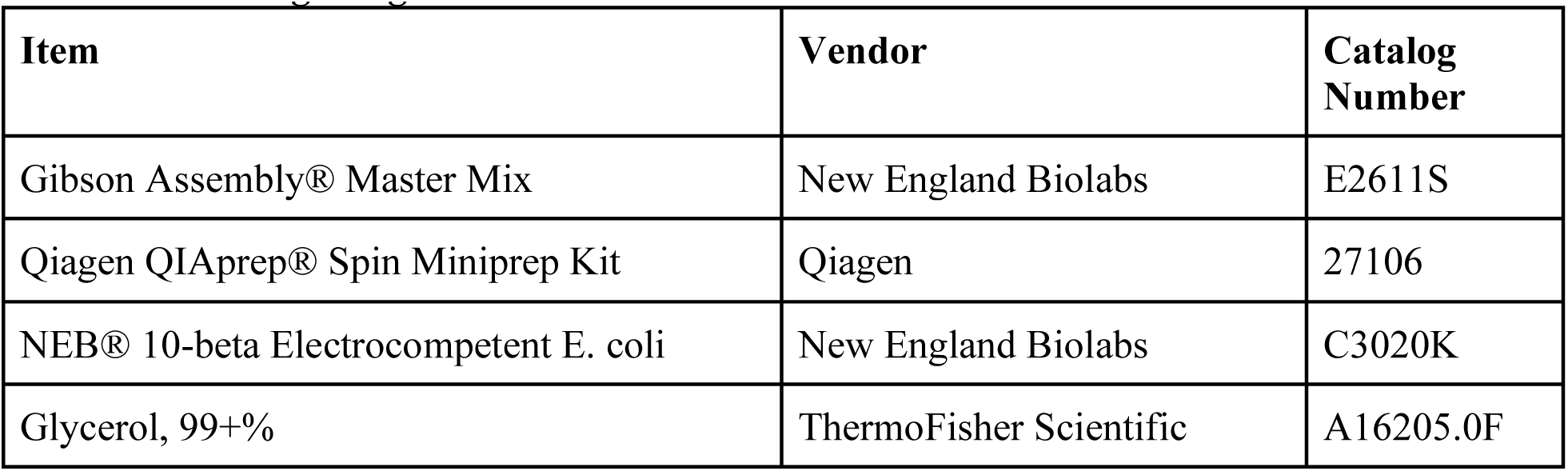
Cloning Reagents.

**Table S6:** Luciferase Reporter Assays Reagents.

| Item | Vendor | Catalog Number |
| --- | --- | --- |
| HEK 293T Cells | ATCC | CRL-3216™ |
| Dulbecco's Modified Eagle's Medium (DMEM), High Glucose | ThermoFisher Scientific | 11965118 |
| Fetal Bovine Serum (Heat Inactivated) | Millipore Sigma (Sigma Aldrich) | F4135-500ML |
| MEM Non-Essential Amino Acids Solution (100X) | ThermoFisher Scientific, Gibco™ | 11140050 |
| GlutaMAX™ Supplement | ThermoFisher Scientific, Gibco™ | 35050061 |
| Sodium Pyruvate (100 mM) | ThermoFisher Scientific, Gibco™ | 11360070 |
| Penicillin-Streptomycin (10,000 U/mL) | ThermoFisher Scientific, Gibco™ | 15140122 |
| Phosphate-buffered saline, no calcium or magnesium (PBS) | ThermoFisher Scientific, Gibco™ | 14190144 |
| Costar® Costar 24-Well Clear TC Treated Plates | Corning® | 3524 |
| FuGENE® HD Transfection Reagent. | Promega | E2311 |
| Opti-MEM™ I Reduced Serum Medium | ThermoFisher Scientific, Gibco™ | 31985088 |
| 24-Well Black Visiplate TC, Case of 14 | Revvity | 1450-605 |
| Thermo Scientific MaxQ 416 High Performance Orbital Shaker | ThermoFisher Scientific | SHKE416HP |
| Dual-Glo® Luciferase Assay System | Promega | E2940 |
| Tecan Infinite® 200 PRO plate reader | Tecan | M Plex |
| pLX304_zeo_mmIrf1 Plasmid | Addgene | 160098 |
| pBMPC3 Bmal1 Plasmid | Addgene | 31367 |
| pCKFB1 Clock Plasmid | Addgene | 31280 |

**Table S7:**
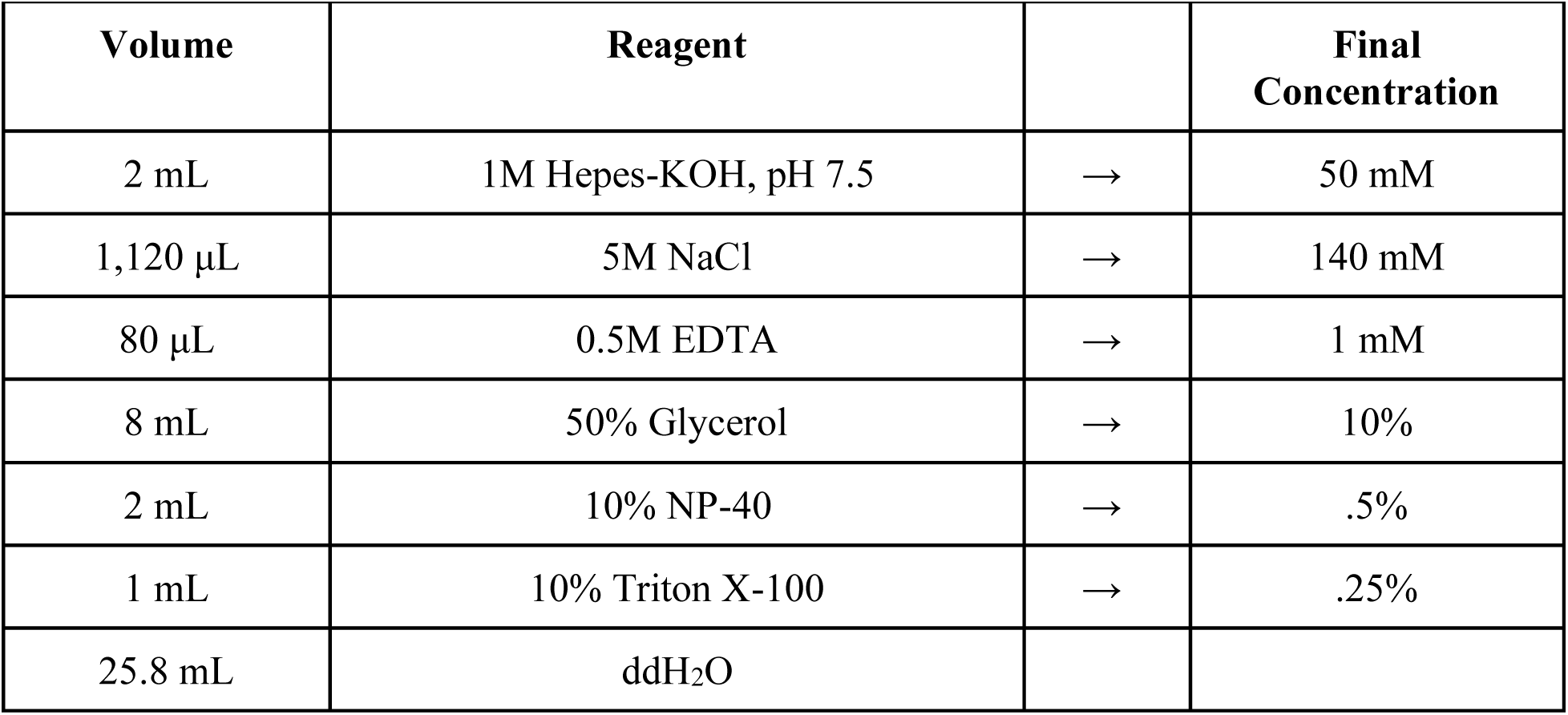
Lysis Buffer 1 (40 mL)

| Volume | Reagent |  | Final Concentration |
| --- | --- | --- | --- |
| 2 mL | 1M Hepes-KOH, pH 7.5 | → | 50 mM |
| 1,120 µL | 5M NaCl | → | 140 mM |
| 80 µL | 0.5M EDTA | → | 1 mM |
| 8 mL | 50% Glycerol | → | 10% |
| 2 mL | 10% NP-40 | → | .5% |
| 1 mL | 10% Triton X-100 | → | .25% |
| 25.8 mL | ddH<sub>2</sub>O |  |  |

**Table S8:** Lysis Buffer 2 (40 mL)

| Volume | Reagent |  | Final Concentration |
| --- | --- | --- | --- |
| 400 µL | 1M Tris-HCl, pH 8.0 | → | 10 mM |
| 1,600 µL | 5M NaCl | → | 200 mM |
| 80 µL | 0.5M EDTA | → | 1 mM |
| 40 µL | 0.5M EGTA | → | 0.5 mL=M |
| 37.8 μL | ddH<sub>2</sub>O | | |

**Table S9:**
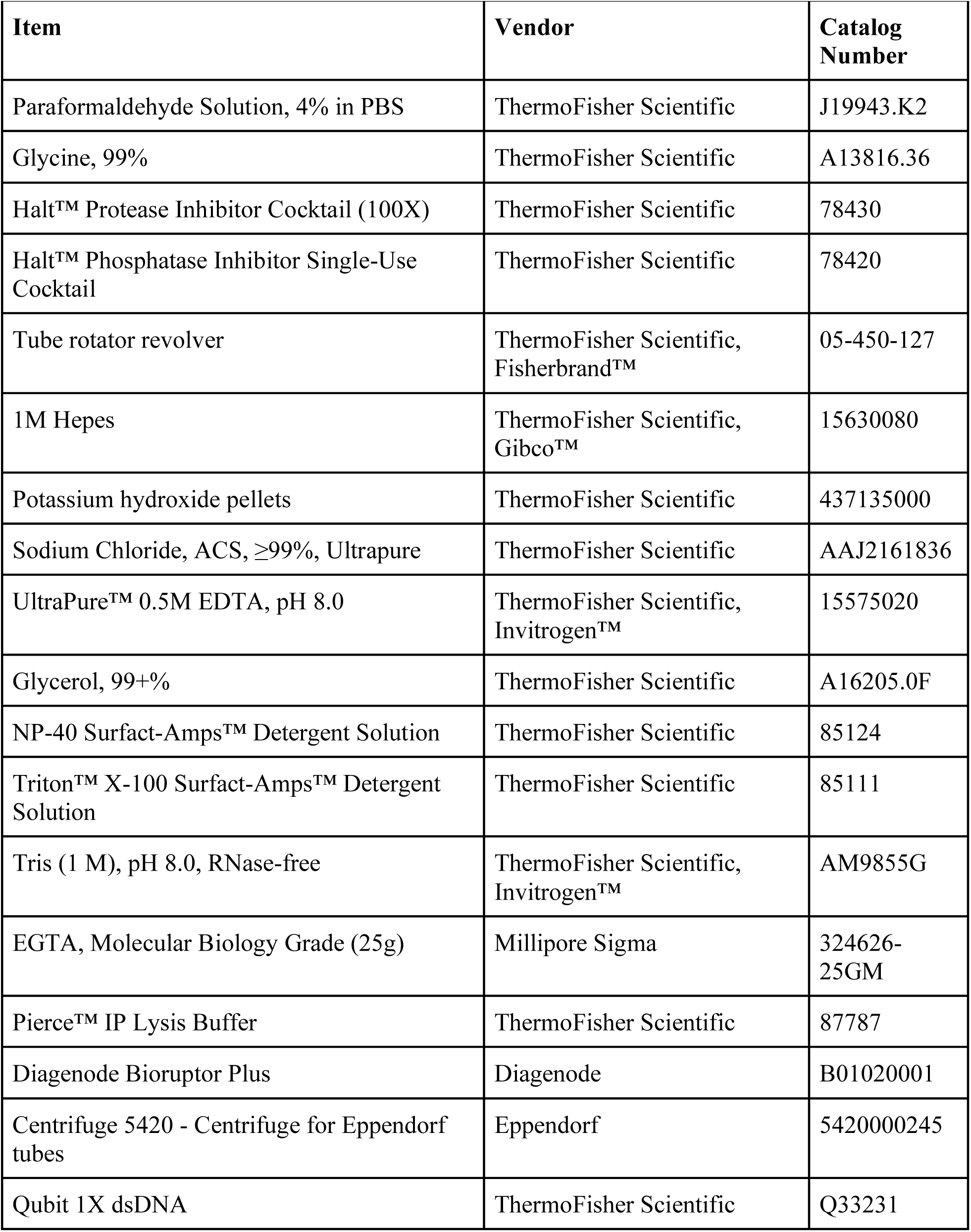
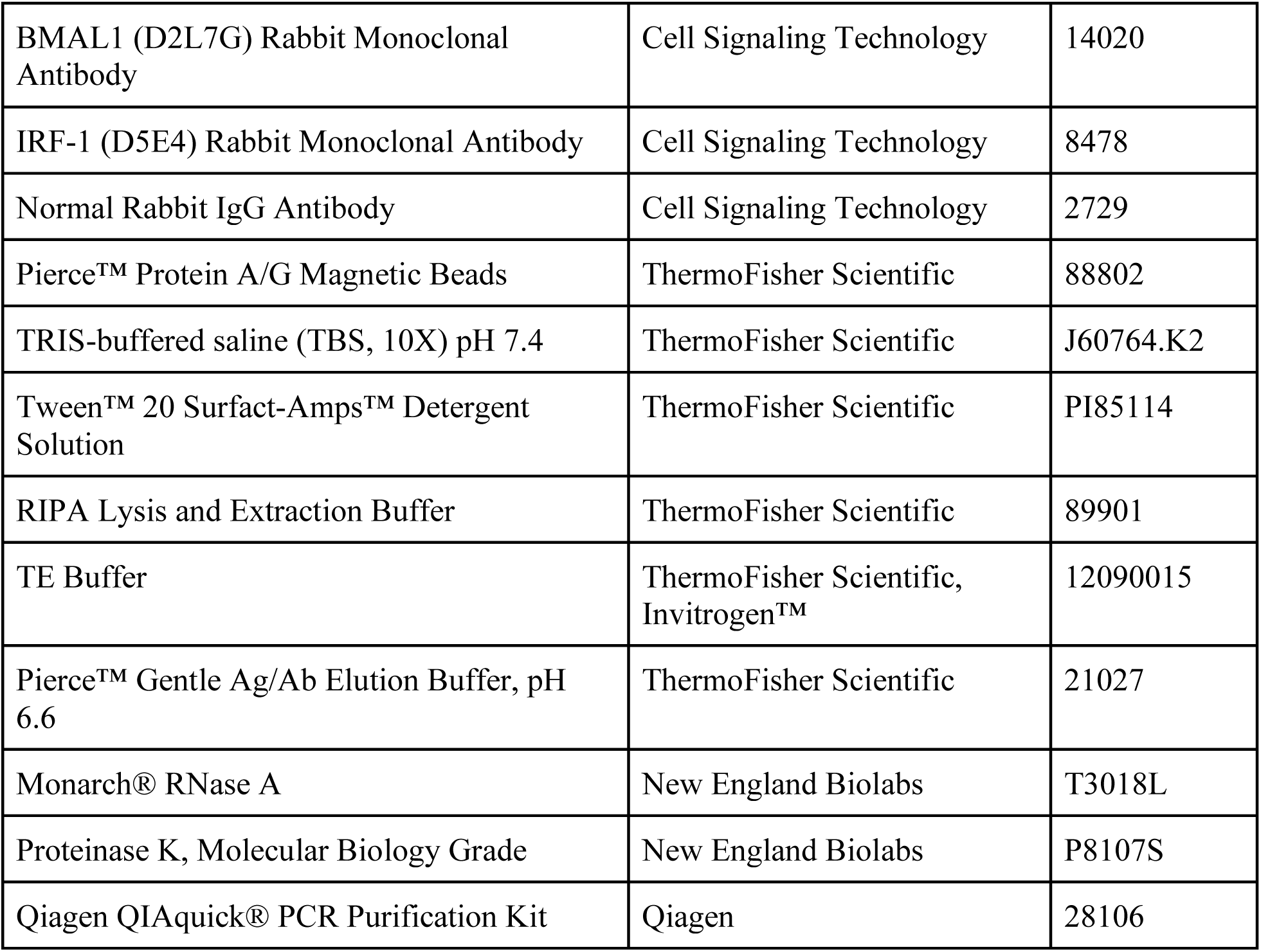
Chromatin Immunoprecipitation (ChIP) Reagents.

| Item | Vendor | Catalog Number |
| --- | --- | --- |
| Paraformaldehyde Solution, 4% in PBS | ThermoFisher Scientific | J19943.K2 |
| Glycine, 99% | ThermoFisher Scientific | A13816.36 |
| Halt™ Protease Inhibitor Cocktail (100X) | ThermoFisher Scientific | 78430 |
| Halt™ Phosphatase Inhibitor Single-Use Cocktail | ThermoFisher Scientific | 78420 |
| Tube rotator revolver | ThermoFisher Scientific, Fisherbrand™ | 05-450-127 |
| 1M Hepes | ThermoFisher Scientific, Gibco™ | 15630080 |
| Potassium hydroxide pellets | ThermoFisher Scientific | 437135000 |
| Sodium Chloride, ACS, ≥99%, Ultrapure | ThermoFisher Scientific | AAJ2161836 |
| UltraPure™ 0.5M EDTA, pH 8.0 | ThermoFisher Scientific, Invitrogen™ | 15575020 |
| Glycerol, 99+% | ThermoFisher Scientific | A16205.0F |
| NP-40 Surfact-Amps™ Detergent Solution | ThermoFisher Scientific | 85124 |
| Triton™ X-100 Surfact-Amps™ Detergent Solution | ThermoFisher Scientific | 85111 |
| Tris (1 M), pH 8.0, RNase-free | ThermoFisher Scientific, Invitrogen™ | AM9855G |
| EGTA, Molecular Biology Grade (25g) | Millipore Sigma | 324626-25GM |
| Pierce™ IP Lysis Buffer | ThermoFisher Scientific | 87787 |
| Diagenode Bioruptor Plus | Diagenode | B01020001 |
| Centrifuge 5420 - Centrifuge for Eppendorf tubes | Eppendorf | 5420000245 |
| Qubit 1X dsDNA | ThermoFisher Scientific | Q33231 |
| BMAL1 (D2L7G) Rabbit Monoclonal Antibody | Cell Signaling Technology | 14020 |
| IRF-1 (D5E4) Rabbit Monoclonal Antibody | Cell Signaling Technology | 8478 |
| Normal Rabbit IgG Antibody | Cell Signaling Technology | 2729 |
| Pierce™ Protein A/G Magnetic Beads | ThermoFisher Scientific | 88802 |
| TRIS-buffered saline (TBS, 10X) pH 7.4 | ThermoFisher Scientific | J60764.K2 |
| Tween™ 20 Surfact-Amps™ Detergent Solution | ThermoFisher Scientific | PI85114 |
| RIPA Lysis and Extraction Buffer | ThermoFisher Scientific | 89901 |
| TE Buffer | ThermoFisher Scientific, Invitrogen™ | 12090015 |
| Pierce™ Gentle Ag/Ab Elution Buffer, pH 6.6 | ThermoFisher Scientific | 21027 |
| Monarch® RNase A | New England Biolabs | T3018L |
| Proteinase K, Molecular Biology Grade | New England Biolabs | P8107S |
| Qiagen QIAquick® PCR Purification Kit | Qiagen | 28106 |

**Table S10:** Western Blot Reagents.

| Item | Vendor | Catalog Number |
| --- | --- | --- |
| T-PER™ Tissue Protein Extraction Reagent | ThermoFisher Scientific | 78510 |
| Pierce™ Protease Inhibitor Tablets, EDTA-free | ThermoFisher Scientific | A32965 |
| Thermo Scientific™ Pierce™ Phosphatase Inhibitor Mini Tablets | ThermoFisher Scientific | A32957 |
| 10x Tris/Glycine/SDS Buffer 1L | Bio-Rad | 1610732 |
| Novex™ Tris-Glycine SDS Sample Buffer (2X) | ThermoFisher Scientific, Invitrogen™ | LC2676 |
| Mini-PROTEAN® TGX™ Protein Gel | Bio-Rad | 4561094 |
| PVDF Transfer Membranes, 0.45 µm, 1 Roll | ThermoFisher Scientific | PI88518 |
| Blotting Grade Blocker Non-Fat Dry Milk | Bio-Rad | 1706404XTU |
| Phosphate-buffered saline, no calcium or magnesium (PBS) | ThermoFisher Scientific, Gibco™ | 14190144 |
| Tween™ 20 Surfact-Amps™ Detergent Solution | ThermoFisher Scientific | PI85114 |
| BMAL1 (D2L7G) Rabbit Monoclonal Antibody | Cell Signaling Technology | 14020 |
| IRF-1 (D5E4) Rabbit Monoclonal Antibody | Cell Signaling Technology | 8478 |
| β-Actin (13E5) Rabbit mAb | Cell Signaling Technology | 4970 |
| OAS2 Antibody | Cell Signaling Technology | 54155 |
| SDHA Monoclonal Antibody | ThermoFisher Scientific, Invitrogen™ | 459200 |
| Anti-rabbit IgG, HRP-linked Antibody | Cell Signaling Technology | 7074S |

**Table S11:** RNA-seq Reagents.

| Item | Vendor | Catalog Number |
| --- | --- | --- |
| TRIzol™ | ThermoFisher Scientific, Invitrogen™ | 15596026 |
| Qiagen RNeasy® Plus Universal Mini Kit | Qiagen | 73404 |
| Qubit™ RNA High Sensitivity (HS) | ThermoFisher Scientific, Invitrogen™ | Q32852 |
| Bioanalyzer High Sensitivity RNA Analysis | Agilent | 5067-1513 |
| Illumina Stranded Total RNA Prep | Illumina | M-GL-02148 v1.0 |
| Illumina RNA Prep with Enrichment | Illumina | M-GL-02145 v1.0 |

**Table S12:** Immunofluorescence Examination Reagents.

| **Item** | **Vendor** | **Catalog Number** |
| --- | --- | --- |
| Bouin's Fixative | ThermoFisher Scientific | 88038 |
| ParaFilm® M Lab Film - 5 Mil, 4" x 125' | Uline | S-25929 |
| Xylene | ThermoFisher Scientific | X3P1GAL |
| Ethanol, Pure, 200 Proof (100%), USP, KOPTEC | Avantor Sciences | 89125-172 |
| Citric acid monohydrate | Millipore Sigma | C1909 |
| Distilled Water | ThermoFisher Scientific | 15230147 |
| Phosphate-buffered saline, no calcium or magnesium (PBS) | ThermoFisher Scientific, Gibco™ | 14190144 |
| BSA | Millipore Sigma | A9418 |
| TRIS-buffered saline (TBS, 10X) pH 7.4 | ThermoFisher Scientific | J60764.K2 |
| Triton™ X-100 Surfact-Amps™ Detergent Solution | ThermoFisher Scientific | 85111 |
| AlexaFluor 488 Donkey anti-rabbit IgG | Abcam | Ab150073 |
| Texas Red Goat Anti-Armenian Hamster IgG | Abcam | Ab5743 |
| DAPI and Hoechst Nucleic Acid Stains Share | ThermoFisher Scientific, Invitrogen™ | D1306 |
| Fluoromount-G ® | ThermoFisher Scientific, Invitrogen™ | 00-4958-02 |
| Prolong Diamond Antifade | ThermoFisher Scientific | P36965 |
| Keyence Fluorescence Microscope BZ-X800 | Keyence | BZ-X800 |

**Table S13:** CSF1R Depletion Experiment Reagents.

| Item | Vendor | Catalog Number |
| --- | --- | --- |
| InVivoMAb anti-mouse CSF1R (CD115) | BioXCell | BE0213 |
| InVivoMAb rat IgG2a isotype control, anti-trinitrophenol | BioXCell | BE0089 |

**Table S14:** Lamina Propria Isolation Buffers.

| Buffer | Reagents |
| --- | --- |
| EDTA-DTT Buffer | 14.25 mL HBSS<br>300 μL of 0.5M EDTA<br>150 μL of 1M DTT<br>300 μL FBS |
| Digest Solution | 4.82 mL RPMI<br>31.25 μL Liberase<br>250 μL DNase I<br>100 μL FBS |
| 80% Percoll | 2 mL Percoll<br>250 μL HBSS<br>250 μL RNase Free H<sub>2</sub>O |

**Table S15:** Lamina Propria Isolation Reagents.

| Item | Vendor | Catalog Number |
| --- | --- | --- |
| HBSS, no calcium, no magnesium, no phenol red | ThermoFisher Scientific, Gibco™ | 14175095 |
| UltraPure™ 0.5M EDTA, pH 8.0 | ThermoFisher Scientific, Invitrogen™ | 15575020 |
| DTT, Crystalline powder | Millipore Sigma | 10197777001 |
| Fetal Bovine Serum (Heat Inactivated) | Millipore Sigma | F4135-500ML |
| RPMI-1640 Medium | Millipore Sigma | R8758 |
| Liberase™ TL Research Grade | Millipore Sigma | 05401020001 |
| DNase I, RNase-free (1 U/μL) Share | ThermoFisher Scientific | EN0521 |
| Cytiva Percoll™ Centrifugation Media | Cytiva | 17089102 |
| TRIzol™ | ThermoFisher Scientific,<br>Invitrogen™ | 15596026 |
| HyClone™ HyPure Water, Molecular<br>Biology Grade | Cytiva | SH30538.FS |
| Phosphate-buffered saline, no calcium or<br>magnesium (PBS) | ThermoFisher Scientific,<br>Gibco™ | 14190144 |

**Table S16:**
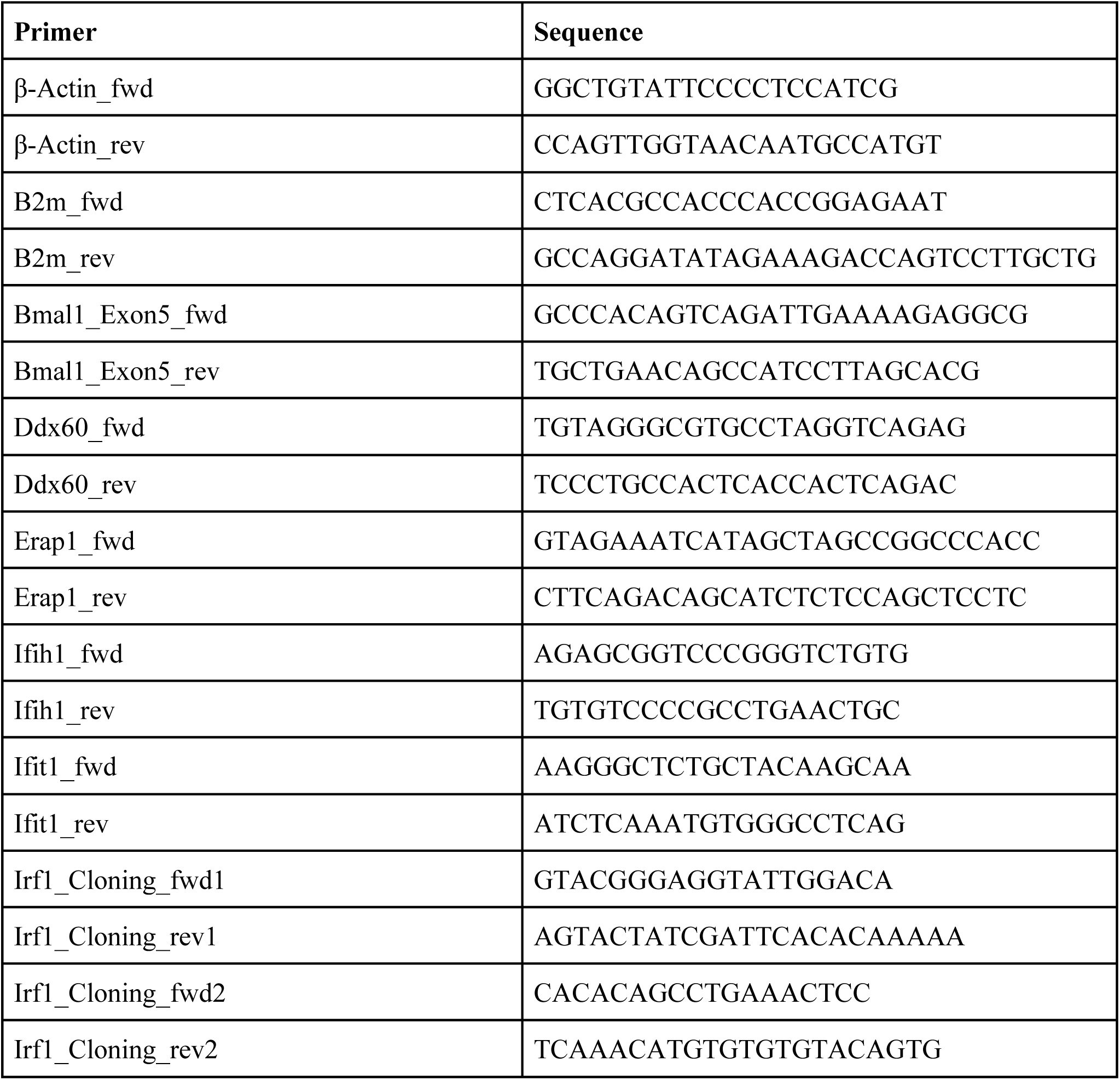
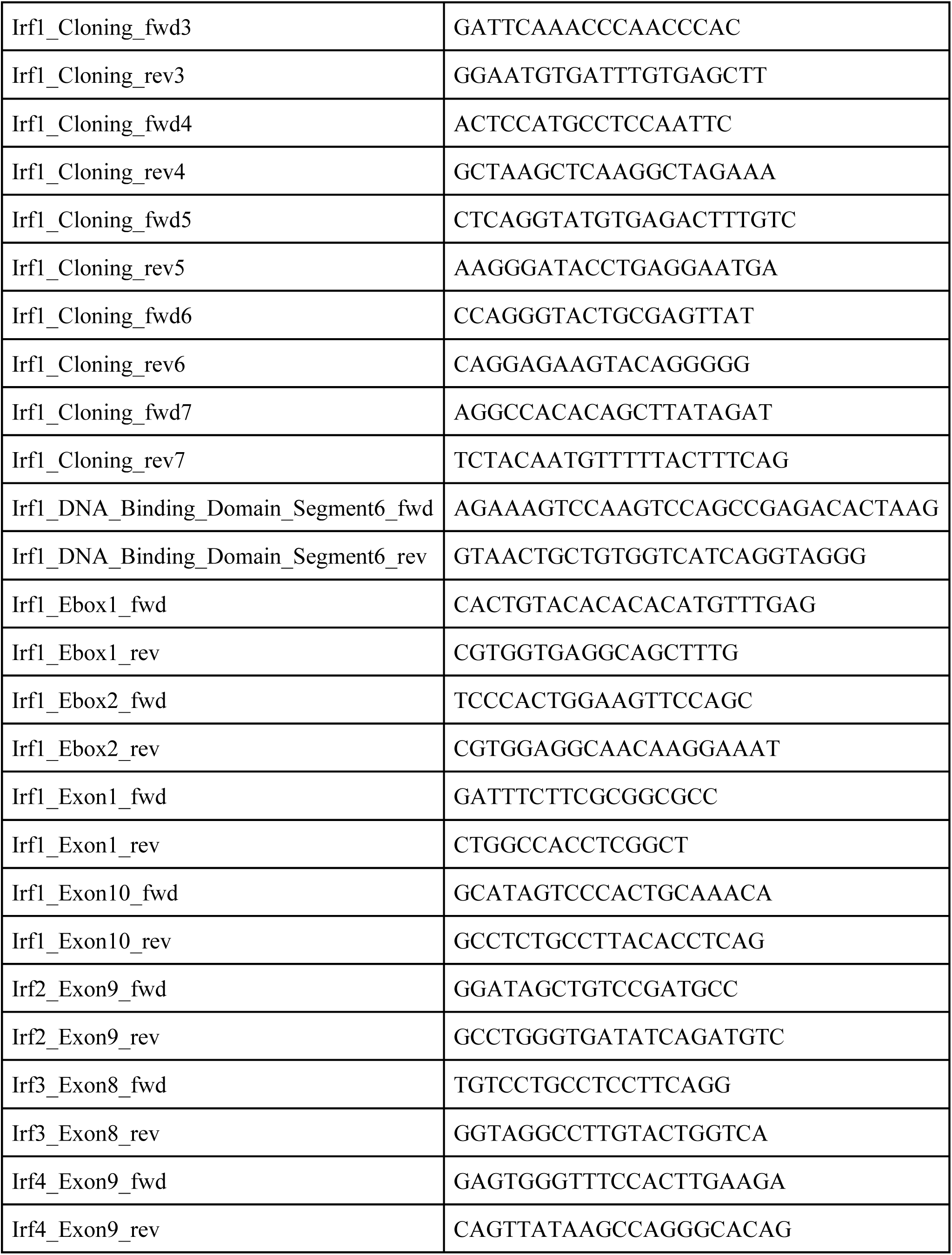
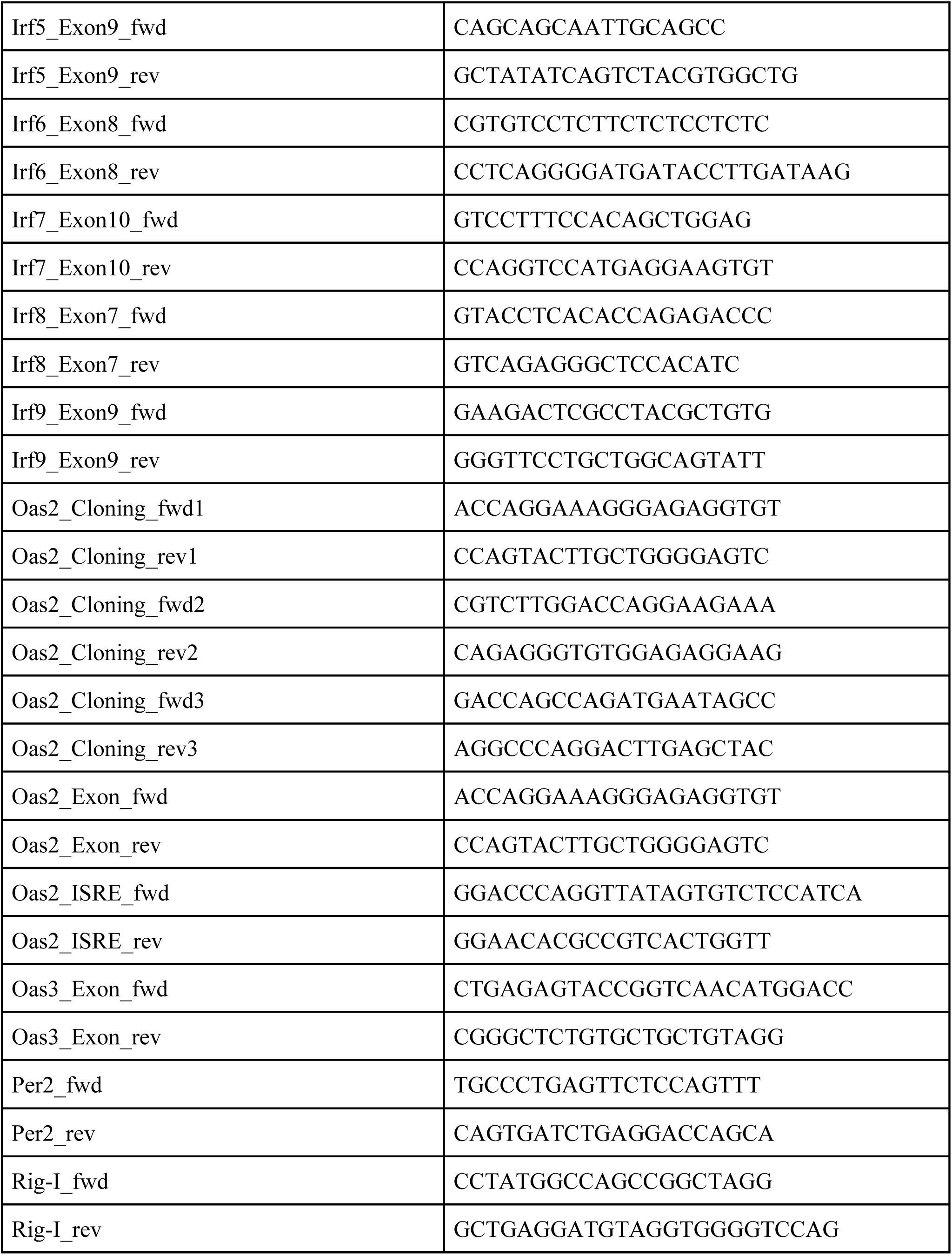
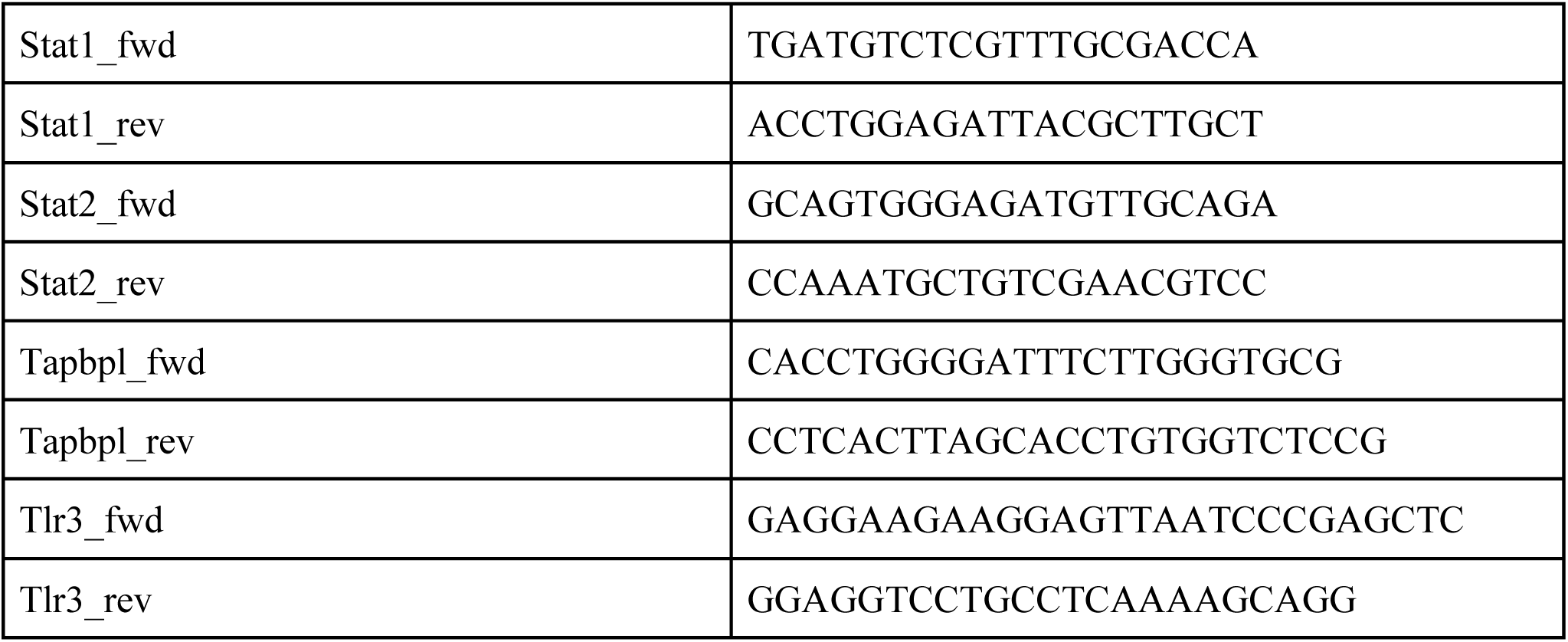
Primers.

